# Single-cell transcriptomics of the early developing mouse cerebral cortex disentangles the spatial and temporal components of neuronal fate acquisition

**DOI:** 10.1101/2020.11.27.401398

**Authors:** Matthieu X. Moreau, Yoann Saillour, Andrzej W. Cwetsch, Alessandra Pierani, Frédéric Causeret

## Abstract

In the developing cerebral cortex, how progenitors that seemingly display limited diversity end up in producing a vast array of neurons remains a puzzling question. The prevailing model that recently emerged suggests that temporal maturation of these progenitors is a key driver in the diversification of the neuronal output. However, temporal constrains are unlikely to account for all diversity across cortical regions, especially in the ventral and lateral domains where neuronal types significantly differ from their dorsal neocortical counterparts born at the same time. In this study, we implemented single-cell RNAseq to sample the diversity of progenitors and neurons along the dorso-ventral axis of the early developing pallium. We first identified neuronal types, mapped them on the tissue and performed genetic tracing to determine their origin. By characterising progenitor diversity, we disentangled the gene expression modules underlying temporal vs spatial regulations of neuronal specification. Finally, we reconstructed the developmental trajectories followed by ventral and dorsal pallial neurons to identify gene waves specific of each lineage. Our data suggest a model by which discrete neuronal fate acquisition from a continuous gradient of progenitors results from the superimposition of spatial information and temporal maturation.

## Introduction

In the nervous system of vertebrates, the production of assorted types of neurons enables the execution of elaborate cognitive functions. Evolutionary changes in neuronal specification processes resulted in the diversification of neuronal types and brain complexification as observed in mammals. In recent years, single-cell RNAseq (scRNAseq) approaches revealed that neuronal diversity is far more extensive than anticipated. For example, the mammalian brain contains more than a hundred excitatory and inhibitory neuron subtypes (Tasic et al., 2018). The developmental mechanisms enabling the emergence of such a diversity are currently under intense investigation.

The neocortex, which has been subjected to strong expansion of cell numbers in primates and especially humans, has attracted most of the attention so far. It is organised in six layers of neurons that are generated according to an “inside-out” sequence (Angevine and Sidman, 1961) and is flanked by three layers cortices: the hippocampus medially and the piriform cortex laterally. All glutamatergic cortical neurons derive from apical progenitors (AP) that reside in the pallial ventricular zone (VZ) and divide to self-renew, produce neurons (direct neurogenesis), or give rise to basal progenitor (BP) displaying a limited proliferation potential in mice (indirect neurogenesis). In the subpallium, ventricular progenitors give rise to inhibitory neurons that will either populate the striatum or migrate dorsally to invade the cortical plate. The so-called pallial-subpallial boundary (PSB) appears initially as a fuzzy border between the dorsal and ventral aspects of the telencephalon that is progressively refined to precisely separate the two compartments (Cocas et al., 2011; Corbin, 2003). These compartments give rise to two building structures of the telencephalon which serve crucial functions at the basis of higher order cognitive and motor behaviours: the cerebral cortex and the basal ganglia. The PSB not only delimitates distinct regions of the developing brain but also acts as a secondary signalling centre, releasing morphogens crucial for cortical patterning (O’Leary and Sahara, 2008). Within the pallium itself, no sharp boundaries between progenitor domains can be found. Instead the graded expression of patterning genes such as *Pax6, Lhx2, Emx1/2* led to the proposal that diversity is not strictly spatially-segregated at the progenitor level as is the case in the developing spinal cord (Pierani and Wassef, 2009; Sagner and Briscoe, 2019; Yun et al., 2001). How the apparent continuity of progenitors gives rise to discrete neuronal types is not clearly established. Currently, temporal regulations are believed to contribute most to neuronal diversity in the cerebral cortex. According to this model, the neurogenic differentiation followed by neocortical neurons is essentially the same and it is the temporal maturation of APs that will influence the final neuronal output (Okamoto et al., 2016; Telley et al., 2019). However, temporal regulations are unlikely to account for all neuronal diversity in the developing pallium, otherwise the hippocampus and piriform cortex would not significantly differ from the neocortex. The tetrapartite model of pallial development (Puelles et al., 1999; Puelles et al., 2000) postulates that AP segregate in four consecutive domains conserved among vertebrates: the medial pallium giving rise to the hippocampus, the dorsal pallium (DP) giving rise to the neocortex, and the lateral and ventral pallium (LP and VP) contributing to the piriform cortex, claustrum and amygdala. However, the precise contribution of LP and VP domains to mature telencephalic structures are still being discussed (Watson and Puelles, 2017; Wullimann, 2017), in part due to complex migratory patterns and to the existence of both laminar and nuclear derivatives. Furthermore, the diversity and identity of LP- and VP-derived neurons remains poorly characterised. To determine the extent of diversity among progenitors and neurons, and apprehend how discrete neuronal types arise from continuous progenitor domains, we implemented a scRNAseq approach. We first characterised neuronal diversity around the PSB, mapped the identified cell types on the tissue and assessed their ontogeny using genetic tracing. We then investigated AP diversity and extensively characterised the molecular signatures of progenitor domains on both sides of the PSB. We used previously published data exploring the temporal maturation of dorsal cortex progenitors (Telley et al., 2019) to discriminate genes whose expression in progenitors changes with the developmental stage from those subjected to spatial regulations along the dorso-ventral (DV) axis. Finally, we reconstructed the developmental trajectories leading to the production of VP- and DP-derived neurons to identify gene waves specific of each lineage. Taken together, our result allow for the first time to unravel the respective contribution of temporal maturation and spatial information to the establishment of neuronal diversity in the developing dorsal telencephalon.

## Material & Methods

### Animals

The following mouse lines were used and maintained on a C57BL/6J background: *Dbx1*^*LacZ*^ (Pierani et al., 2001), *Emx1*^*Cre*^ (Gorski et al., 2002), *ΔNp73*^*Cre*^ (Tissir et al., 2009), *Wnt3a*^*Cre*^ (Yoshida et al., 2006), *Rosa26*^*YFP*^ (Srinivas et al., 2001) and *Rosa26*^*tdTomato*^ (Madisen et al., 2010). All animals were handled in strict accordance with good animal practice as defined by the national animal welfare bodies, and all mouse work was approved by the French Ministry of Higher Education, Research and Innovation as well as the Animal Experimentation Ethical Committee of Paris Descartes University (CEEA-34, licence number: 18011-2018012612027541).

### scRNAseq

Four wild-type E12.5 embryos obtained from two distinct litters were collected in ice-cold HBSS. A region spanning from the subpallium to the DP (see Fig. 1A) was dissected on both hemispheres and dissociated using the Neural Tissue Dissociation Kit (P) (Miltenyi Biotec) and a gentleMACS Octo Dissociator with Heaters following the manufacturer’s instructions. Cell clumps and debris were removed by two consecutive rounds of centrifugation 3 minutes at 200xg and filtration through 30 µm cell strainers (Miltenyi Biotec). Additional mechanical dissociation and removal of cell aggregates was achieved by gentle pipetting up and down 5 times using a P200 pipette and Gel Saver tips (QSP), followed by filtration through a 10 µm cell strainer (pluriSelect). Assessment of cell viability and counting were achieved using a MACSQuant flow cytometer (Miltenyi Biotec) in the presence of 7-AAD. The absence of dead cells, debris or doublets in the cell suspension was crosschecked using a hemocytometer and trypan blue staining. Approximatively 10,000 cells were loaded on a 10X Genomics Chromium Controller. The entire procedure, from the sacrifice of the pregnant female until loading the controller, was achieved in two hours. A single-cell barcoded cDNA library was prepared using a Chromium Single Cell 3’ Library and Gel Bead Kit v2 and sequenced at a total depth of 190 million reads on an Illumina NextSeq500 sequencer. Raw sequencing reads were processed to counts matrix with Cell Ranger (version 2.2.1) with default parameters, using the mm10 mouse genome as reference. All statistical analyses were performed under R (version 3.6.3).

**Figure 1.**
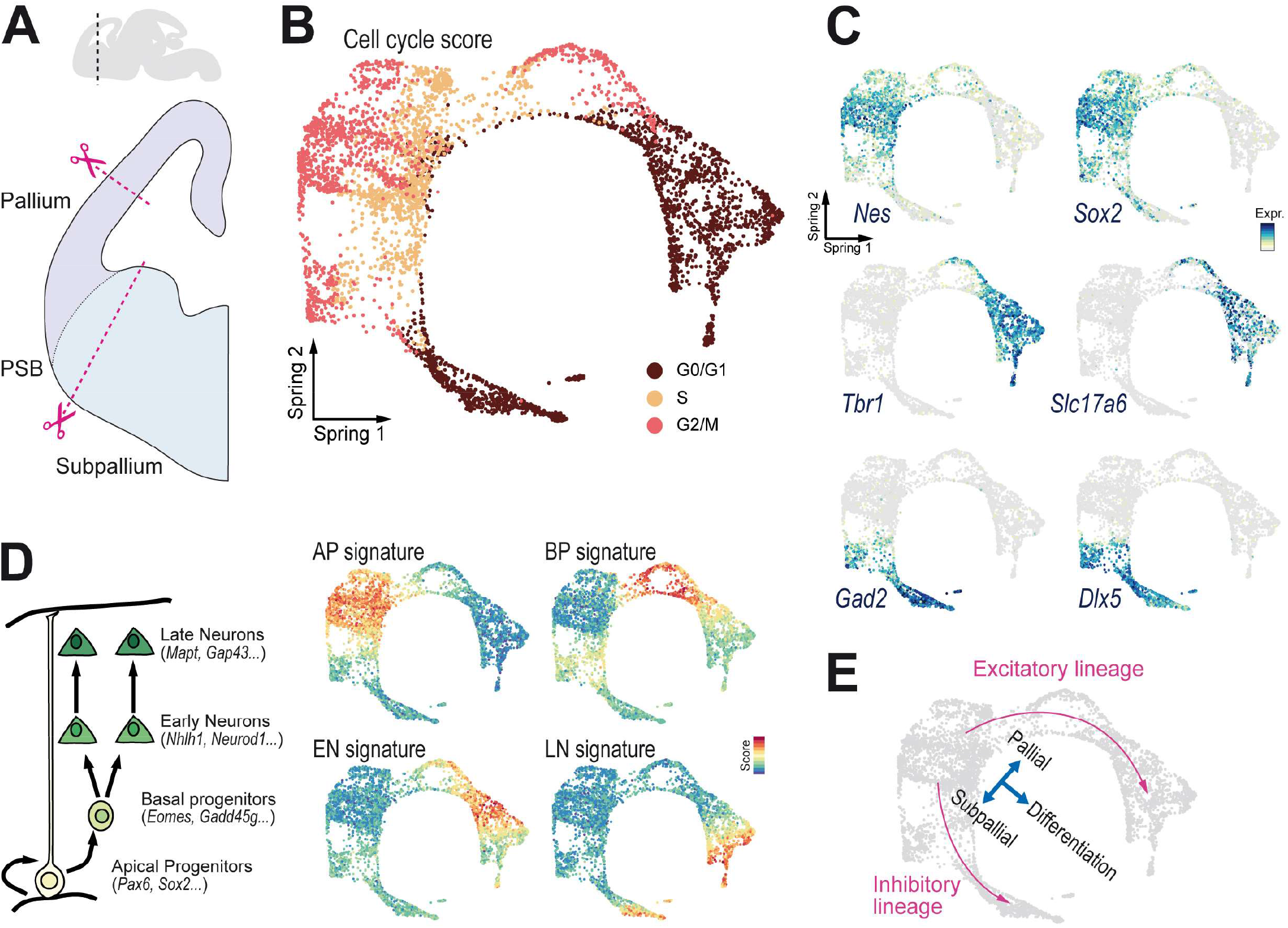
Sampling cellular diversity around the PSB. **A**. The region encompassing the pallial-subpallial boundary (PSB) was dissected from E12.5 mouse embryos and subjected to scRNAseq. **B**. SPRING representation of the dataset with cells (points) coloured according to their cell cycle state. **C**. Expression of genes defining neural progenitors (*Nes* and *Sox2*), glutamatergic neurons (*Tbr1* and *Slc17a6*/Vglut2) and GABAergic neurons (*Gad2* and *Dlx5*). **D**. SPRING visualisation of the signature scores for apical progenitors (AP), basal progenitors (BP), early neurons (EN) and late neurons (LN). **E**. Summary of the dataset topology, a continuity of differentiating excitatory and inhibitory neurons emerges from a group of progenitors. The main axes of the SPRING representation correspond to the differentiation state (progenitors vs neurons) and the region of origin (pallium vs subpallium).

### Cell filtering

To retain only high quality cells, we applied filters on unique molecular identifier (UMI), gene and mitochondrial counts per cell. We first filtered cells based on the percentage of UMI associated with mitochondrial transcripts and number of detected genes and retained cells within a 3 median absolute deviation (MAD) around the population median for these metrics. We further excluded cells having a number of UMI above 3 MADs of the population median. Potential doublets were removed using Scrublet (Wolock et al., 2019). Finally, we excluded genes detected in less than 3 cells. Dimensionality reduction was performed using the SPRING tool (Weinreb et al., 2018). After dimensionality reduction, 11 cells sharing neighbours between excitatory and inhibitory neurons, and therefore likely corresponding to undetected doublets, were removed manually. The raw counts value for the 4225 high quality cells was library size normalised and scaled using Seurat (v 2.3.4) (Butler et al., 2018).

### Cell state score computation

We reasoned that clustering procedure gives meaningful results only if applied to cell populations in the same broad transcriptional state (e.g. apical progenitors or late neurons). To select cells in the same transcriptional state we computed apical progenitors (AP), basal progenitors (BP), early neurons (EN) and late neurons (LN) signature scores using lists of published and manually selected genes (Telley et al., 2016; the detailed list is available in the annotated codes, see Data availability below) as input to the Seurat function ‘addModuleScore’. The scores were differentially combined to define and extract the populations of interest by K-means clustering prior to downstream analysis.

### Neuronal populations clustering

To explore neuronal transcriptional diversity we first applied iterative graph-based clustering using scrattch.hicat (v0.0.16) (Tasic et al., 2016) on glutamatergic and GABAergic neurons separately after removing cell cycle, ribosomal or mitochondrial associated genes form the input count matrix. At each clustering iteration step we further excluded principal components (PCs) highly correlating with sequencing depth, percentage of ribosomal or mitochondrial transcripts detected per cells.

Because variation among some glutamatergic neuron clusters were obviously associated with different maturation states we further excluded those with a LN score < 1. Hierarchical grouping of the remaining clusters was performed with the scrattch.hicat function ‘build_dend’ using the mean gene expression profile across clusters as input. To defined clusters specific transcriptional signatures we further subset the clusters core cells as assigned more than 95/100 trials to the same cluster using the centroid classifier of the ‘map_sampling’ function. Top cluster marker genes were identified using a receiver operating characteristic (ROC) analysis implemented in the Seurat function ‘FindMarkers’. Barplot representation of gene expression was built using the scrattch.vis package (v0.0.210).

To identify modules of genes coexpressed across different neuronal clusters we implemented the WGCNA package (v1.68) (Langfelder and Horvath, 2008). We used the normalised expression matrix of the first 1000 most variable genes defined with the Seurat function ‘FindVariableGenes’ as input and a softpower = 4 to compute the signed adjacency matrix. Modules were identified by clustering over the distance obtained from the topological overlap matrix (TOM) and their expression profile were summarised as their module eigengene.

### Apical progenitors analysis

We aimed at ordering APs by performing a new SPRING dimensionality reduction while excluding cell cycle associated genes. Cells were projected along the main axis of variation by fitting a principal curve using princurve (v2.1.4) as described previously (Mayer et al., 2018). Genes differentially expressed along the inferred DV axis were identified using the ‘differentialGeneTest’ function of the Monocle package (v2.14) (Trapnell et al., 2014). Progenitor cells were grouped in 7 clusters using partition around medoids on the smoothed expression profile of the significantly differentially expressed genes. Genes with similar expression dynamics were grouped by hierarchical clustering. To predict temporal maturation scores for apical progenitors we took advantage of the Telley et al., (2019) dataset. After retrieving the raw count matrix from the GEO repository (GSE118953) we extracted cells sorted 1h after FlashTag injection. Based on their expression profile, we excluded medial pallium APs as well as putative doublets with GABAergic interneurons to retain 725 high quality cells (E12: 178 cells, E13: 202 cells, E14: 124 cells, E15: 221 cells). We performed principal component analysis (PCA) dimensionality reduction using the Seurat function ‘RunPCA’ over all differentially expressed genes between the distinct sampling groups identified with the Wilcoxon rank sum test implemented in the ‘FindAllMarkers’ function. Cells were projected onto a pseudo-maturation axis by fitting a principal curve over the first 6 PCs. We trained a random-forest regression model implemented in the Caret package (v6.0-84) to predict the cells’ pseudo-maturation score (position along the axis) using the scaled expression value of the top 1129 variable genes selected with the ‘FindVariableGenes’ function. The 100 most important features were set as input to re-optimise the regression model which was used to predict temporal maturation for APs in our dataset. To investigate the spatial and temporal components we first identified genes with variable expression in pallial apical progenitors along the DV axis using the ‘differentialGeneTest’ function as described above, and those differentially expressed between E12 and E15 1h after FlashTag injection in the Telley et al. dataset. The “temporal module” was defined as the intersection between the two sets of differentially expressed genes. We then computed the Pearson’s correlation coefficient between the smoothed expression matrices over 200 points of temporal genes in our dataset and that of Telley et al. Genes of the temporal module showing coordinated changes in expression in the two datasets were grouped using hierarchical clustering. The ‘spatial module’ was defined as genes differentially expressed along the DV axis at E12 but which do not show variation between E12 and E15 in the Telley et al. dataset.

### Differentiation trajectories inference

In order to predict fate bias along the pallial differentiation path we performed FateID (v0.1.9) (Herman et al., 2018) analysis setting mature pallial neuron clusters (at the exception of CRs) as “attractor” states. We focused on the two trajectories that were traced back to APs for pseudotime ordering. We restricted our analysis to the confidently assigned cells by selecting those biased towards one lineage with > 50% of votes and an absolute difference of votes between the two lineages > 25% (Nr4a2 lineage: 617 cells, Bhlhe22 lineage: 520 cells). Cells were projected along a pseudotime axis by fitting a principal curve on the SPRING dimensionality reduction that captures the variation associated with pan-neuronal differentiation. We tested for differential gene expression along pseudotime between the two trajectories using the likelihood ratio test implemented in Monocle’s function ‘differentialGeneTest’. Trajectory specific enrichment was determined by adapting the Area Between Curves (ABC) method from Monocle. Finally, genes with similar smoothed trends where clustered using partition around medoids.

To identify genes best predicting early transcriptional differences between the two trajectories at the BP state, we perform ROC analysis as implemented in Seurat’s function ‘FindMarkers’ and use the Area Under Curve (AUC) value to rank genes according to their ability to classify BP cells in their respective lineages.

### Data availability

Raw and processed data can be retrieved from the GEO database (accession number GSM4910731). Comprehensive and annotated R codes used in this study can be found at https://matthieuxmoreau.github.io/EarlyPallialNeurogenesis/

### Tissue processing

For staging of the embryos, midday of the vaginal plug was considered as embryonic day 0.5 (E0.5). Embryos were collected in ice-cold PBS, dissected, and immediately fixed by immersion in 4% paraformaldehyde, phosphate buffer (PB) 0.12 M, pH7.4 for 2 hours at 4°C. Samples were cryoprotected by incubation in 10% sucrose, PB overnight at 4°C, embedded in 7.5% gelatine, 10% sucrose, PB and frozen by immersion in isopentane cooled at −55°C. 20 µm thick sections were obtained with a Leica CM3050 cryostat and collected on Superfrost Plus slides (Menzell-Glasser).

### Immunostaining

The following primary antibodies were used: chick anti-GFP (Aves Labs GFP-1020, 1:1000), chick anti-βGal (Abcam ab9361, 1:1000), rabbit anti-Nfib (Atlas Antibodies HPA003956, 1:1000), rabbit anti-Foxp2 (Abcam ab16046, 1:1000), goat anti-Nurr1 (R&D systems AF2156, 1:200), mouse anti-Meis2 (Sigma WH0004212M1, 1:1000) and guinea pig anti-Bhlhb5 (Skaggs et al., 2011, 1:100). The following secondary antibodies were obtained from Jackson ImmunoResearch: donkey anti-chick Alexa-488 (1:1000), donkey anti-mouse Cy5 (1:500), donkey anti-guinea pig Cy5 (1:500), donkey anti-rabbit Cy3 (1:700), donkey anti-rabbit Cy5 (1:500), donkey anti-goat Cy3 (1:700) and donkey anti-goat Cy5 (1:500). DAPI (1µg/mL) was used for nuclear staining. Slides were mounted in Vectashield (Vector).

### In situ Hybridisation

For each gene of interest, a DNA fragment (typically 500-800bp) was amplified from an embryonic brain cDNA library using Phusion polymerase (Thermo) and a pair of specific primers (Table S1). The promoter sequence of the T7 RNA polymerase (GGTAATACGACTCACTATAGGG) was added in 5’ of the reverse primer. Alternatively, for *Dbx1, Emx1, Gsx2, Reln, Rorb, Sp8* and *Trp73*, a plasmid containing part of the cDNA was linearised by enzymatic restriction. Antisense digoxigenin-labelled RNA probes were then obtained by in vitro transcription using T7 RNA polymerase (New England Biolabs) and digRNA labelling mix (Roche). In situ hybridisation was carried out as previously described (Schaeren-Wiemers and Gerfin-Moser, 1993) using a hybridisation buffer composed of 50% formamide, 5X SSC, 1X Denhardt’s, 10% dextran sulfate, 0.5 mg/mL yeast RNA, 0.25 mg/mL herring sperm DNA. Probes were detected using an anti-digoxigenin antibody coupled to alkaline phosphatase (Roche) and NBT/BCIP (Roche) as substrates. Slides were mounted in Mowiol.

### Image acquisition

In situ hybridisation images were obtained using a Hamamatsu Nanozoomer 2.0 slide scanner. Immunofluorescence images were acquired using a Leica SP8 confocal microscope with a 40X objective. Quantifications were achieved by counting the proportion of GFP^+^ cells among the marker-positive population using the ImageJ software. Data are represented as mean ± sd, individual values are plotted.

## Results

### Sampling cellular diversity around the developing ventral pallium

Using a droplet-based approach (10x Genomics Chromium V2 platform), we performed unbiased scRNAseq on dissected explants containing the PSB and adjacent tissue, obtained from four wild-type E12.5 embryos (Fig. 1A). After quality control and filtering, we obtained 4225 cells sequenced at a median depth of 11,665 UMI and a median detection of 3,571 genes per cell.

Visualisation of transcriptional heterogeneity on a SPRING plot – a 2D projection using force-directed layout of the k-nearest-neighbor (Weinreb et al., 2018) – revealed that the dataset is organised in a “crab-shaped” topology consisting in a large group of cycling cells from which stemmed two branches of postmitotic (G0/G1 phase) cells (Fig. 1B). Expression of pallial (*Tbr1*), excitatory neurons (*Slc17a6*/Vglut2) and inhibitory neurons markers (*Dlx5* and *Gad2/*GAD65) revealed that the two branches correspond to pallial excitatory and subpallial inhibitory differentiation trajectories, respectively (Fig. 1C). Consistently, the large group of cycling cells was found to express *Sox2* and *Nes*/Nestin, genes characteristic of neural progenitors (Fig. 1C).

To ensure that we successfully captured all states through which pallial cells transit from progenitor to mature neuron, we calculated expression scores for apical (AP) or intermediate/basal (BP) progenitor and neurons at early (EN) or late (LN) stages of differentiation using sets of genes previously shown to characterise these distinct steps (Telley et al., 2016). Visualisation of the scores on the SPRING plot embedding recapitulates the known sequence of transcriptional states by which cells transit from AP to LN, indicating that we sampled the different intermediates along the differentiation trajectory of pallial neurons (Fig. 1D).

When focusing on cells from the glutamatergic branch with high BP signature score (Fig. S1), we noticed that one group expressing genes associated with proliferation such as *Top2a, Mki67, Cdk1* or *Aurka*, and annotated as S or G2/M phase cells, appears to diverge from another group of aligned G0/G1 cells which do not express such cell cycle genes (Fig. 1B, S1). The latter most likely corresponds to the trajectory followed by cells transitioning from AP to neurons without achieving a last division as BP, hence representing a non-proliferative intermediate state. This observation suggest that at E12, direct neurogenesis involves transition through a “BP-like” transcriptomic state because we could not see any evidence for existence of trajectories skipping this intermediate state. If such a skipping trajectory was existing between AP and EN state, we would expect to observe cells expressing transcript from both identities, which we did not find. We therefore concluded that, at least at E12, progenitors undergoing direct neurogenesis transit through a BP-like state *Eomes*^+^ (Tbr2), *Btg2*^+^ (Tis21) and *Neurog2*^+^ without replicating their DNA or dividing.

Taken together, these results indicate that the two main axes in the low-dimensional manifold of our dataset reflect (i) the differentiation status, from cycling progenitor to postmitotic mature neuron and (ii) the pallial or subpallial origin of the cells (Fig. 1E). Thus our data recapitulate global tissue organization around the PSB at E12.5.

### Molecular and spatial characterization of neuronal populations

To explore neuronal diversity, we first selected mature neurons based on their high LN and low EN scores using K-means clustering (Fig. 2A). We then performed iterative graph-based clustering using the scrattch.hicat pipeline from the Allen institute (Tasic et al., 2016) to identify glutamatergic and GABAergic cell types which, expectedly, formed two distinct groups (Fig. 2B).

**Figure 2.**
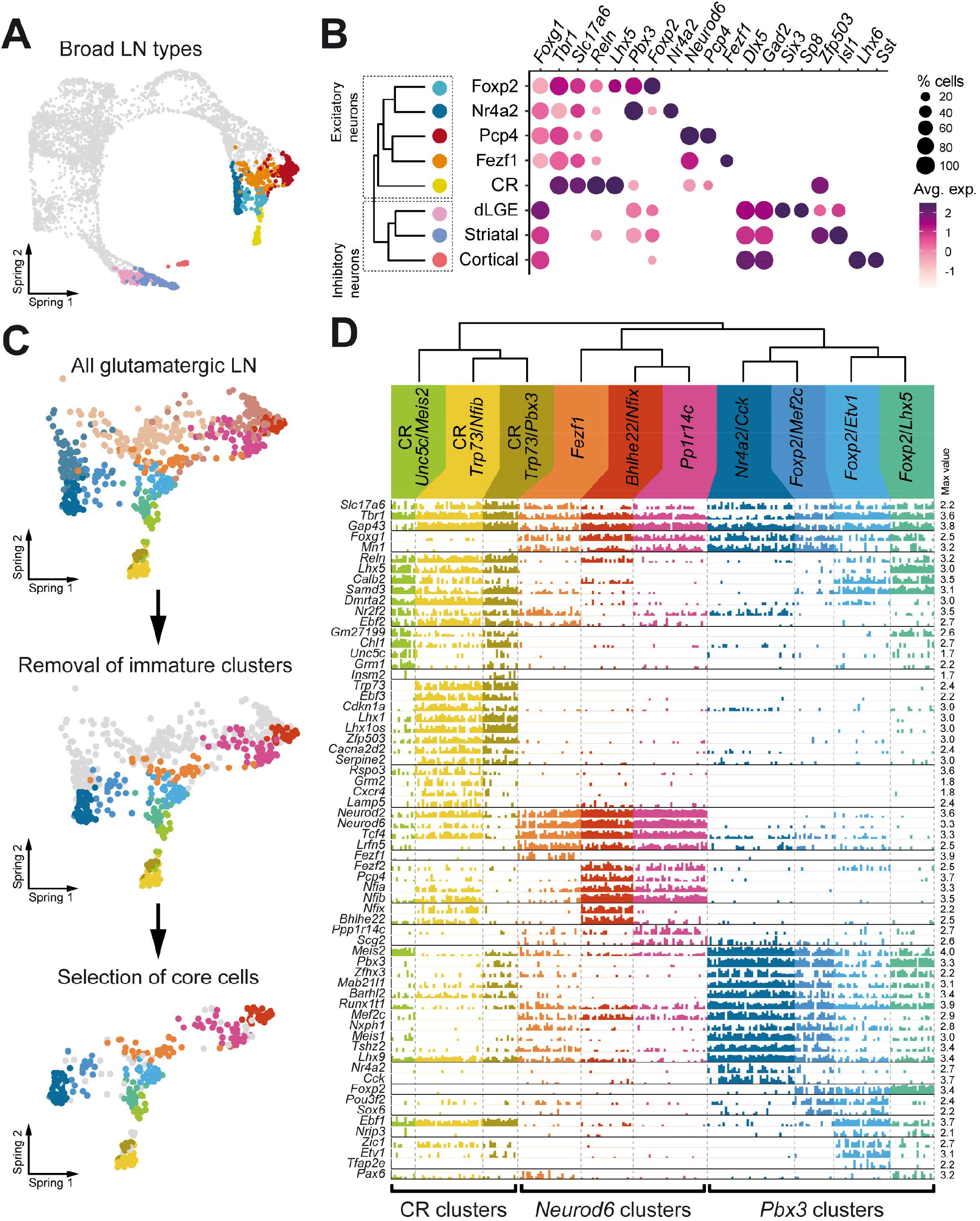
Neuronal diversity around the PSB. **A**. SPRING visualisation of single cells coloured by broad neuronal types (5 glutamatergic and 3 GABAergic). **B**. Bubble chart representing the expression of genes that allowed to identify the broad neuronal types. Circle size represents the proportion of cells expressing the gene considered in each cluster, the average expression level is color-coded. Hierarchical relation between cell clusters is depicted by the dendrogram. **C**. Strategy to define clusters that confidently reflect existing mature excitatory neuron populations. Clusters composed of immature cells were removed as well as cells that were not systematically attributed to the same cluster using centroid classifier. **D**. Bar plot representing the expression level of selected genes differentially expressed between clusters which allow to finely define their identity. Each bar corresponds to a single cell, bars height corresponds to the expression level of a given gene in a given cell. The dendrogram on the top indicates the hierarchical relation between cell clusters.

Three main GABAergic subtypes were identified (Fig. 2B) and could be further subdivided (Fig. S2). A first group displayed the characteristic *Lhx6*/*Sst*/*Calb1* signature of cortical interneurons migrating from the medial ganglionic eminence and preoptic area. Accordingly, this cluster appeared disconnected from the main inhibitory differentiation trajectory on the SPRING plot (Fig. S2A), suggesting that the progenitors and differentiation intermediates were not captured by the dissection. Two clusters contained neurons expressing the dorsal lateral ganglionic eminence (dLGE) marker gene *Sp8* (Ma et al., 2012), and could be split by that of either *Six3* or *Zic1* (Fig. S2) The remaining clusters were characterised by the strong expression of *Isl1, Ebf1* and *Zfp503* (Nolz1), genes reported to be expressed by striatal projection neurons (Chang et al., 2004; Corbin, 2003; Garel et al., 1999). Within glutamatergic neurons, analysis of the higher hierarchical relationship revealed 3 major groups (Fig. 2B) corresponding to Cajal-Retzius (CR) cells, the only *Foxg1*-negative cells (Hanashima, 2004), branching apart from *Foxg1*^*+*^ cells which were split into two main groups, with on one side *Neurod6*^+^ neurons, and on the other side those expressing *Pbx3* (Fig. 2B). To accurately characterise glutamatergic neurons we first decided to exclude clusters with a median LN score < 1 to retain only the most mature type (Fig. 2C). This approach lead to the identification of 10 mature glutamatergic neuronal types, for which we defined the transcriptional signature using only clusters’ core cells, i.e. those assigned more than 95 out of 100 trials to the same cluster using a centroid classifier (Figs 2C, 2D, S3). The genes significantly enriched in each cluster are listed in Table S2.

### Cajal-Retzius cells subtypes

CRs have been recently reported to form a transcriptionally distinct group of neurons in the cortex (Tasic et al., 2018; Yao and Tasic, 2020). Accordingly, hierarchical clustering indicated that CR clusters diverge from all other glutamatergic neurons (Fig. 2B,D). They were readily identified by the absence of *Foxg1* (Hanashima, 2004) but also *Mn1*, and the expression of previously described markers such as *Reln, Lhx5, Calb2* (Calretinin), *Dmrta2* (Dmrt5), *Nr2f2* (COUP-TFII) and *Ebf2* (Chuang et al., 2011; Miquelajauregui et al., 2010; Ogawa et al., 1995; Saulnier et al., 2013; Tripodi, 2004) (Figs 2D, S3B). Of note, none of these genes is fully specific to CRs, most noticeably *Reln* was found in both *Bhlhe22*^+^ and *Foxp2*^*+*^ populations, although at lower levels than in CRs (Fig. 2D).

Hierarchical clustering indicated that CRs divide in two main groups: the first one expressing a set of genes among which *Trp73, Ebf3, Cdkn1a, Lhx1, Cacna2d2* and a smaller one displaying higher, albeit not restricted, expression of *Unc5c, Grm1* (mGluR1/Lot1) and high levels of *Calb2* and *Meis2* (Figs 2D, S3I). CRs originate from several distinct regions of the forebrain and *Trp73* expression is a well-described feature of those derived from medial sources i.e. the cortical hem, pallial septum and thalamic eminence (Griveau et al., 2010; Meyer et al., 2002; Ruiz-Reig et al., 2017; Tissir et al., 2009), whereas VP-derived CRs where previously shown to be p73-negative and Calretinin^high^ (Griveau et al., 2010; Hanashima et al., 2007). We therefore concluded that the first hierarchical distinction between CRs subtypes is that regarding their medial vs VP origin.

Among *Trp73*^+^ CRs, a large cluster displayed a strong signature consisting of known genes expressed in CRs such as *Grm2* (mGluR2) or *Cxcr4* (Stumm et al., 2003; Yamazaki et al., 2004) combined with dorsal cortex markers (*Nfib, Nfix*) (Plachez et al., 2008), whereas a smaller cluster had a *Insm2*^*+*^*/Pbx3*^*+*^*/Mab21l1*^*+*^ signature and shared the expression of *Chl1* and *Grm1* with VP-derived CRs (Figs 2D, S3J).

In situ hybridisation patterns (Fig. 3A,C) indicated that *Trp73*-negative CRs (VP-derived) are mostly confined to the lateral cortex and lateral olfactory tract (LOT) region, consistent with their low *Nfib* expression and previous reports (Bielle et al., 2005). Regarding p73^+^ CRs (those from medial sources), the *Insm2*^*+*^*/Pbx3*^*+*^ cells were found around the LOT and in the subpallial marginal zone (MZ) whereas *Grm2*^*+*^*/Nfib*^*+*^ subtypes were found above the entire dorsal cortex (Fig. 3A,C). These data suggest that the *Grm2*^*+*^ population corresponds to hem-derived CR and the *Insm2*^*+*^ to septum and/or thalamic eminence-derived CRs.

**Figure 3.**
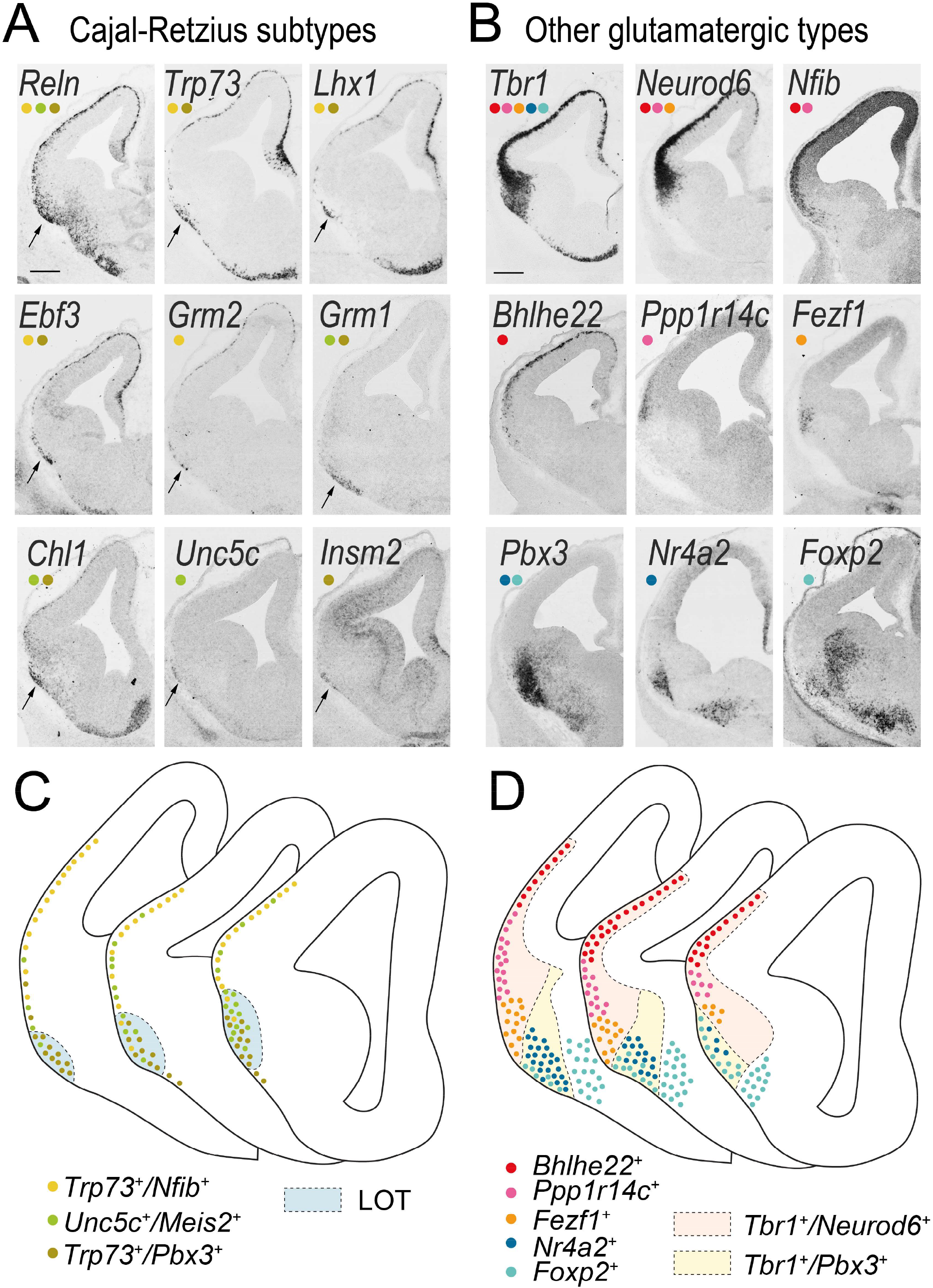
Mapping neuronal populations on tissue. **A, B**. In situ hybridisation of selected genes expressed specifically in Cajal-Retzius cells clusters (A) or other glutamatergic clusters (B) on E12.5 forebrain coronal sections. Coloured dots indicate the neuronal populations expressing the gene considered. **C, D**. Schematic representation of the position occupied by Cajal-Retzius cells subtypes (C) and other glutamatergic subtypes (D). Scale bar: 200 µm.

We performed fate-mapping experiments in order to assess whether the two p73^+^ CR subpopulations correspond to distinct origins. Using the *Wnt3a*^*Cre*^ line to label hem-derived CRs, we found all of them to be Nfib-positive, including those that migrated down close to the LOT region (Fig. 4A). By contrast, the *ΔNp73*^*Cre*^ line, which labels CRs from the hem, septum and thalamic eminence, allowed to identify some Nfib-negative CRs at all medio-lateral levels (Fig. 4B). We also found Meis2-positive cells especially around the LOT and occasionally in the dorsal pallium (Fig. 4C). We therefore concluded that the *Trp73*^*+*^*/Nfib*^*+*^ subset indeed corresponds to hem-derived CRs whereas the *Trp73*^*+*^*/Pbx3*^*+*^ cells most likely derive from the septum and/or thalamic eminence. Our data are therefore consistent with previous histological characterization of CR subtypes and demonstrate that VP-derived CRs clearly diverge from other subtypes generated at medial sources.

**Figure 4.**
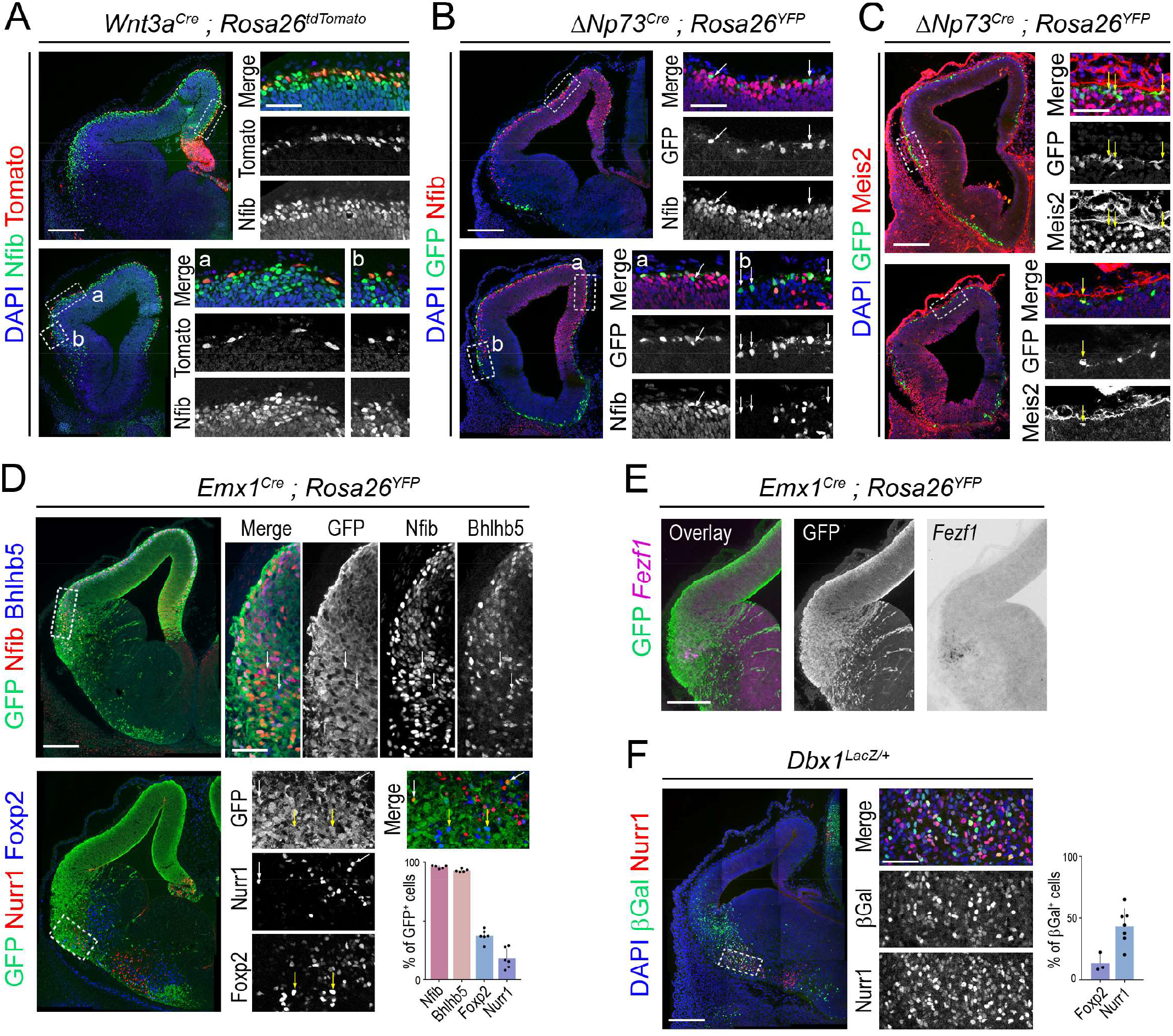
Genetic tracing of excitatory neuronal subtypes. **A**. Immunostaining of E12.5 coronal sections from *Wnt3a*^*Cre*^;*Rosa26*^*tdTomato*^ embryos with an antibody against Nfib. Tomato-positive cells were all found to express Nfib, regardless of the region considered. **B**. E12.5 coronal sections from Δ*Np73*^*Cre*^;*Rosa26*^*YFP*^ embryos stained for Nfib and GFP. Some GFP-positive cells were found to be Nfib-negative (arrows). **C**. Immunostaining of E12.5 coronal sections from Δ*Np73*^*Cre*^;*Rosa26*^*YFP*^ embryos with antibodies against Meis2 and GFP. Some Meis2-positive cells were found among GFP+ cells (arrows), mostly around the lateral olfactory tract. **D**. Immunostaining of E12.5 coronal sections from *Emx1*^*Cre*^;*Rosa26*^*YFP*^ embryos with antibodies against GFP, Nfib, Bhlhb5, Nurr1 and Foxp2. In the top panel, white arrows point to some of the rare GFP-negative Nfib+/Bhlhb5+ cells. In the bottom panel, white arrows point to the few Nurr1+ cells which are GFP-positive and yellow arrows to Foxp2+ cells which are GFP-positive. The histogram (mean ± sd) indicates the proportion of GFP+ cells among those expressing the given markers. Individual measurements are plotted, each dot corresponds to one section. **E**. Overlay between serial sections from the same E12.5 *Emx1*^*Cre*^;*Rosa26*^*YFP*^ embryo stained either with a GFP antibody or a *Fezf1* probe. **F**. E12.5 coronal section from *Dbx1*^*LacZ/+*^ embryos with antibodies against βGal and Nurr1. The histogram (mean ± sd) indicates the proportion of βGal+ cells among those expressing Nurr1 but also Foxp2. Scale bars: 200 µm for low magnification images and 50 µm for high magnification panels.

### Neurod6 clusters

Among the *Neurod2*^*+*^*/Neurod6*^*+*^/*Tcf4*^*+*^ clusters (node G in Fig. S3A), the first population to branch apart was characterised by the specific expression of the transcription factor coding gene *Fezf1* (Figs 2D). In situ hybridisation for *Fezf1* stained a sharp domain in the ventral aspect of the prospective insular cortex (Fig. 3B,D). The remaining two clusters shared the expression of *Nfia, Nfib, Pcp4* and *Fezf2* and were distinguished by that of either *Nfix*/*Bhlhe22* (Bhlhb5) or *Ppp1r14c/Scg2* (Figs 2D, S3H). In situ hybridisation revealed that *Bhlhe22* and *Nfix* are expressed in the preplate of the dorsal pallium whereas *Ppp1r14c* and *Scg2* are found more laterally (Figs 3B,D, S4). The three *Neurod6*^*+*^ populations therefore occupy distinct dorsoventral positions. We then used genetic tracing in the *Emx1*^*Cre*^ line, that recombines mostly in LP and DP progenitors (Gorski et al., 2002), to find out that 96±1% of Nfib^+^ and 93±1% of Bhlhb5^+^ cells derive from Emx1 lineages and are therefore LP/DP derivatives, consistent with their position (Fig. 4D). We also found that the in situ hybridisation signal for *Fezf1* overlaps with the ventral-most GFP^+^ area in *Emx1*^*Cre*^;*Rosa26*^*YFP*^ embryos suggesting that this population also derives from the LP/DP (Fig. 4E). However, in the absence of LP or DP-specific Cre driver, and despite clear distinctions in the relative dorso-ventral position of *Bhlhe22*^+^, *Ppp1r14c*^+^ *and Fezf1*^*+*^ populations, it is difficult to ascertain whether they can be classified as DP or LP derivatives.

### Pbx3 clusters

Four populations were found among the *Pbx3*/*Mab21l1*^*+*^ branch (Figs 2D). Three of them showed expression of the Forkhead box protein 2 transcription factor *Foxp2*, which is also expressed by striatal neurons outside of the *Tbr1*^+^ domain (Fig. 2B). In situ hybridisation pattern for the *Foxp2* transcript indicate that these populations are located around the LOT all along the antero-posterior axis (Fig. 3B,D). The three populations could be distinguished by the expression of either *Lhx5, Mef2c* or *Etv1* (Er81), although these genes are also detected in other clusters (Figs 2D, S3). Tissue mapping indicated that *Etv1*^*+*^ cells are mostly found in rostral regions (Fig. S4, Zimmer et al., 2010). By contrast, the *Lhx5*^+^ population is distributed more caudally and could correspond to neurons migrating from the thalamic eminence towards the olfactory bulb (Huilgol et al., 2013). The *Foxp2*/*Mef2c* population could not be precisely mapped due to the absence of specific marker. The last cluster corresponded to neurons specifically expressing the genes *Nr4a2*, coding for the orphan nuclear receptor Nurr1, and *Cck* coding for the peptide hormone Cholecystokinin (Fig. 2D). Among Tbr1-positive pallial derivatives, these cells occupy the most ventral position (Fig. 3B,D). Interestingly these neurons also showed the lowest *Tbr1* but highest *Lhx9* expression (Fig. 2D), which is the signature of the previously described VP migrating stream (Tole, 2005). To determine the origin of *Nr4a2*^+^ and *Foxp2*^+^ neurons, we took advantage of the specific expression of the transcription factor Dbx1 in VP progenitors (Bielle et al., 2005; Medina et al., 2004) to perform lineage tracing experiment and compare with *Emx1*^*Cre*^;*Rosa26*^*YFP*^ embryos. We favoured short-term lineage tracing using *Dbx1*^*LacZ/+*^ embryos to permanent tracing with *Dbx1*^*Cre*^ as a delay in recombination was previously shown to prevent labelling of the earliest Dbx1-derived cells (Bielle et al., 2005; Puelles et al., 2016b). We found that a higher proportion of Foxp2 immunoreactive cells derive from the Emx1 lineage (37±5%) than the Dbx1 lineage (14±7%). Conversely, Nurr1^+^ cells are more often Dbx1-derived (43±14%) than Emx1-derived (18±9%). We concluded that *Nr4a2*^+^ neurons are mostly VP-derived whereas *Foxp2*^+^ are mostly LP-derived although care should be taken in the interpretation of these data as (i) β–Galactosidase expression is not permanent in *Dbx1*^*LacZ/+*^ animals, which may lead to underestimate the contribution of Dbx1-expressing progenitors, and (ii) a certain extent of recombination in the VP is evident in the *Emx1*^*Cre*^ line (sometimes even in subpallium, see Fig. 4D, E) and we cannot formally exclude postmitotic recombination in VP derivatives. Nevertheless, *Nr4a2*^+^ neurons clearly appear as the most ventral pallial glutamatergic type in our dataset. Our data therefore indicate that neuronal identities in the LP and VP are not sharply distinguished by tracing from Emx1- or Dbx1-expressing progenitor.

### Spatial identity of neuronal subtypes

*Pbx3*^*+*^ and *Neurod6*^+^ populations occupy distinct and complementary pallial domains. This could be imposed by the different lineages they belong to and/or by the position cells eventually occupy. As a first step in distinguishing between these possibilities, we reasoned that if regionally imposed identity exists it should be shared by neurons from very different identities but occupying the same histological location. This implies that we should be able to find gene modules common to GABAergic subtypes and excitatory neurons localised nearby the PSB. We investigated this hypothesis using weighted gene correlation network analysis (WGCNA) (Langfelder and Horvath, 2008) to identify co-expressed gene modules. By applying this analysis to all late neurons present in our dataset, we identified 6 co-expression modules (Fig. S5 and Table S3). Analysis of the expression profile of each module (represented as their eigengene) indicated that two of them are restricted to either GABAergic or glutamatergic lineages and a third one to CRs. These modules therefore reflect neuronal identity (Fig. S5A). Two additional modules displayed inverted gradients of expression between DP and VP-derived excitatory neurons. They could therefore depict the lineage of origin, the spatial identity, or both. Finally, the last module eigengene was found at high levels in VP-derived neurons but also among GABAergic cells (especially dLGE neurons), correlating with the spatial positioning of these cells rather than any other feature. Interestingly, this latter module was found to segregate the hem-derived CR cluster (*Trp73*/*Nfib*) that reside in the dorsal cortex, from the two smaller CR clusters that occupy mostly ventral positions on the tissue (Fig. S5B). A reverse segregation of CR clusters was observed using the DP eigengene. Our data therefore support the existence of gene modules not only linked to lineage-related features but also reflecting spatial identity as imposed by the position of cells along the dorso-ventral axis.

### Diversity of Radial Glia progenitors is best described by continuous gradients of gene expression

In order to determine how neuronal diversity emerges, we focused our attention on APs. The diversity of APs within and among dorso-ventral domains of the pallium remains poorly characterised. To better understand the extent of differences between VP, LP and DP progenitors, we first extracted APs by K-means clustering based on the signature scores previously calculated (Fig. 1D). We then decided to align cells according to their estimated position on a pseudo DV axis to see whether boundaries would emerge. To this end, we performed a SPRING dimensionality reduction after excluding the cell-cycle associated genes, and subsequently projected cells on the principal curve capturing the main source of transcriptional variability among progenitors (Fig. 5A). Finally, AP distribution was discretised using partition around medoids over the smoothed expression of the significantly differentially expressed genes along this axis. We found that dividing APs in seven groups (three in the subpallium and four in the pallium) was the best trade-off allowing to correctly position the PSB without overclusterizing the data (Fig. 5B,C). On the basis of gene expression, pallial APs were found to split into one VP cluster, two LP clusters, and one DP cluster. Genes with differential expression along the pseudo DV axis are represented in Figs 5C and S6, and listed in Table S4. Our approach was validated by the observation that known genes such as *Gsx2, Dlx1/2* or *Olig2* were restricted to subpallial clusters, *Dbx1* expression was detected almost exclusively in the VP domain (even though its expression is weaker in APs than BPs) and *Emx1* was found absent from subpallial clusters and steadily increased with the pseudo DV score (Figs 5C,D, S6). Consistently the expression dynamics of *Ascl1, Pax6, Sp8* and *Emx2* was found graded along the pseudo DV axis, matching the known expression patterns of these genes (Fig. S6). As expected, the PSB constituted the most remarkable border among APs. We found that it is best defined by the expression of *Gsx2* on the subpallial side and *Tfap2c* or *Dmrta2* on the pallial side (Figs 5D,E, S6) consistent with previous findings (Desmaris et al., 2018; Konno et al., 2019). We identified multiple additional genes with sharp PSB expression border, including novel ones such as *Gm29260* or *Lypd6* (Figs 5D,E, S6). However, the overwhelming majority of genes differentially expressed along the pseudo DV axis did not display sharp borders. For example, *Celf4, Sema5a* or *Fat4* peak at the VP but are also found in adjacent domains (Fig. S6). *Sfrp2* which is often used as a PSB marker, is detected at low levels in all domains with the few cells apposed to the PSB showing highest expression (Fig. S6).

**Figure 5.**
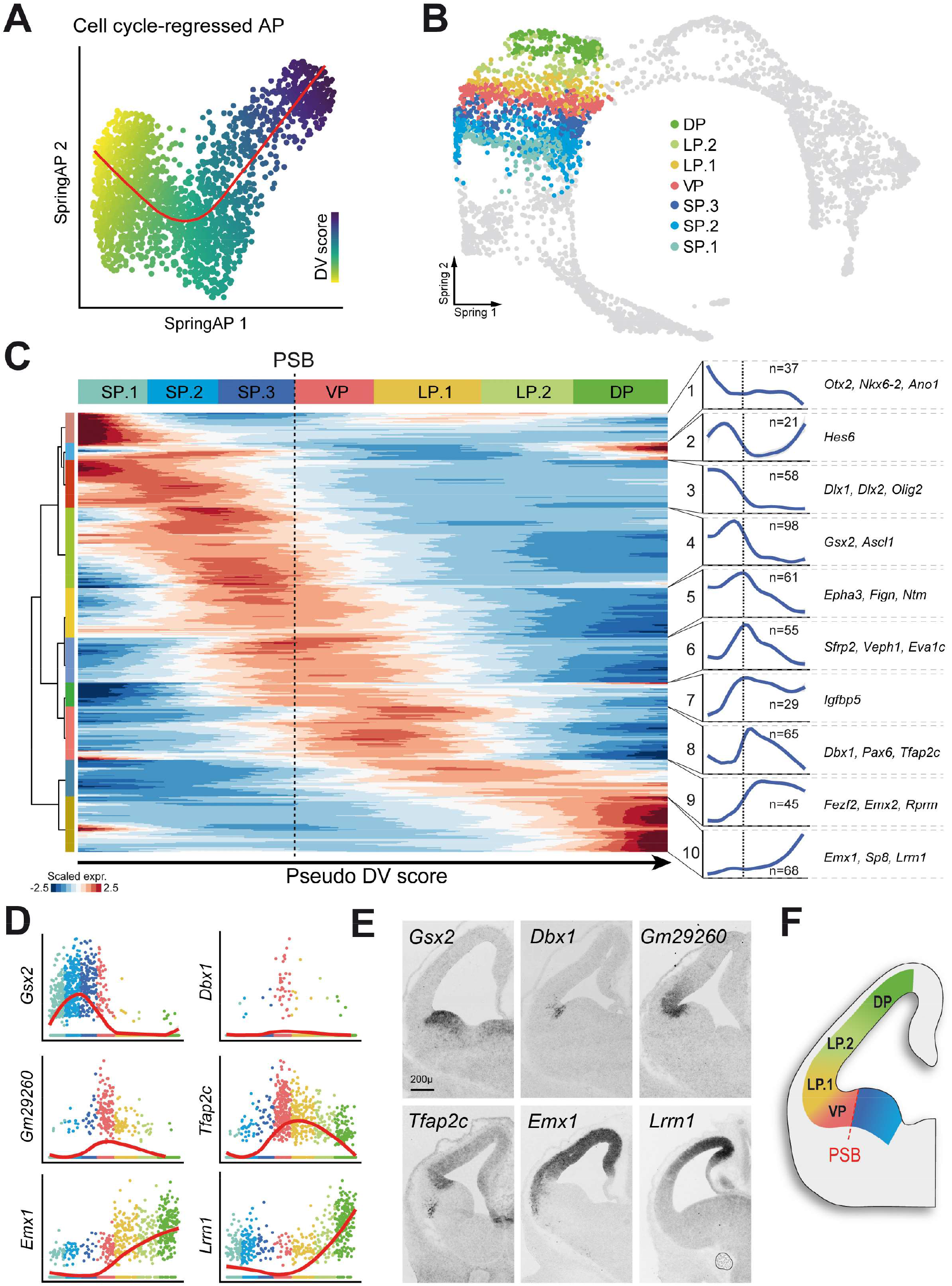
Diversity of apical progenitors. **A**. SPRING embedding of apical progenitors (AP) after excluding cell-cycle associated genes. The red line indicates the principal curve on which cells are projected to determine a pseudo dorso-ventral (DV) score. **B**. SPRING visualisation of single cells coloured according to AP clusters. **C**. Heatmap illustrating the expression of the 537 variable genes along the pseudo DV axis. The dashed line indicates the PSB. Genes with similar patterns were grouped in clusters (indicated by numbers) whose average expression trend is represented by the blue curves on the right **D**. Expression level of selected genes in APs along the pseudo DV position. Each dot corresponds to one single AP color-coded according to the domain it belongs to. The red curve indicates the smoothed expression profile. **E**. In situ hybridisation on sections of wild-type E12.5 embryos for the same genes as D. **F**. Drawing of the estimated position of the AP clusters characterised. Scale bar: 200 µm.

Since our approach only focused on differences along the DV axis, we also evaluated if other sources of variations could account for the diversity of pallial progenitors. We thus subjected pallial APs to iterative graph-based clustering while excluding all sources of variation correlating with either the inferred DV position or the cell cycle score. This allowed us to identify one group of APs characterised by the expression of a handful of genes such as *Id3* or *Fgf15*, that were found enriched in the caudal part of the VP by in situ hybridisation (Fig. S6C). Beyond this domain, we were unable to find evidence of major differences in the transcriptional identity of APs along the antero-posterior axis of the pallium. Overall, we concluded that genes with sharp expression on either side of the PSB coexist with genes that encompass this boundary, indicating that a certain degree of continuity in APs occurs even at the PSB. By contrast, within the pallium the superimposition of genes with graded expression in APs seems to be the rule.

Our data therefore provide a high resolution atlas of gene expression along the DV axis of the early developing telencephalic VZ and supports the hypothesis that besides the few genes that sharply delineate the PSB, progenitor identity changes gradually according to the position of cells.

### Early spatial identity within pallial APs reflects temporal dorsal-pallial maturation

The cerebral cortex displays a well-known neurogenic gradient in which neurogenesis is initiated in ventral regions and progressively extends to dorsal territories (Bayer and Altman, 1991). It was recently shown that neuronal diversity in the developing neocortex arises mostly from the temporal progression of APs (Okamoto et al., 2016; Telley et al., 2019). We therefore queried in which extent the distinction between LP, VP and DP progenitors could mirror that of young vs old dorsal cortex APs.

To test such a possibility, we used the time-series dataset from Telley et al. (2019), which describes the transcriptomic signature of E12 to E15 APs from the dorsal pallium, and trained a regression model to predict a maturation score for each AP present in our dataset (Fig. 6A). We found clear difference in the maturation estimate between clusters as the VP displayed the highest predicted score whereas the DP had the lowest score (Fig. 6B). In other words, VP progenitors display a more mature transcriptomic profile than their DP counterparts. This also indicates that part of the DV axis transcriptional differences can be explained by the expression of genes responsible for the temporal maturation of dorsal cortex APs.

**Figure 6.**
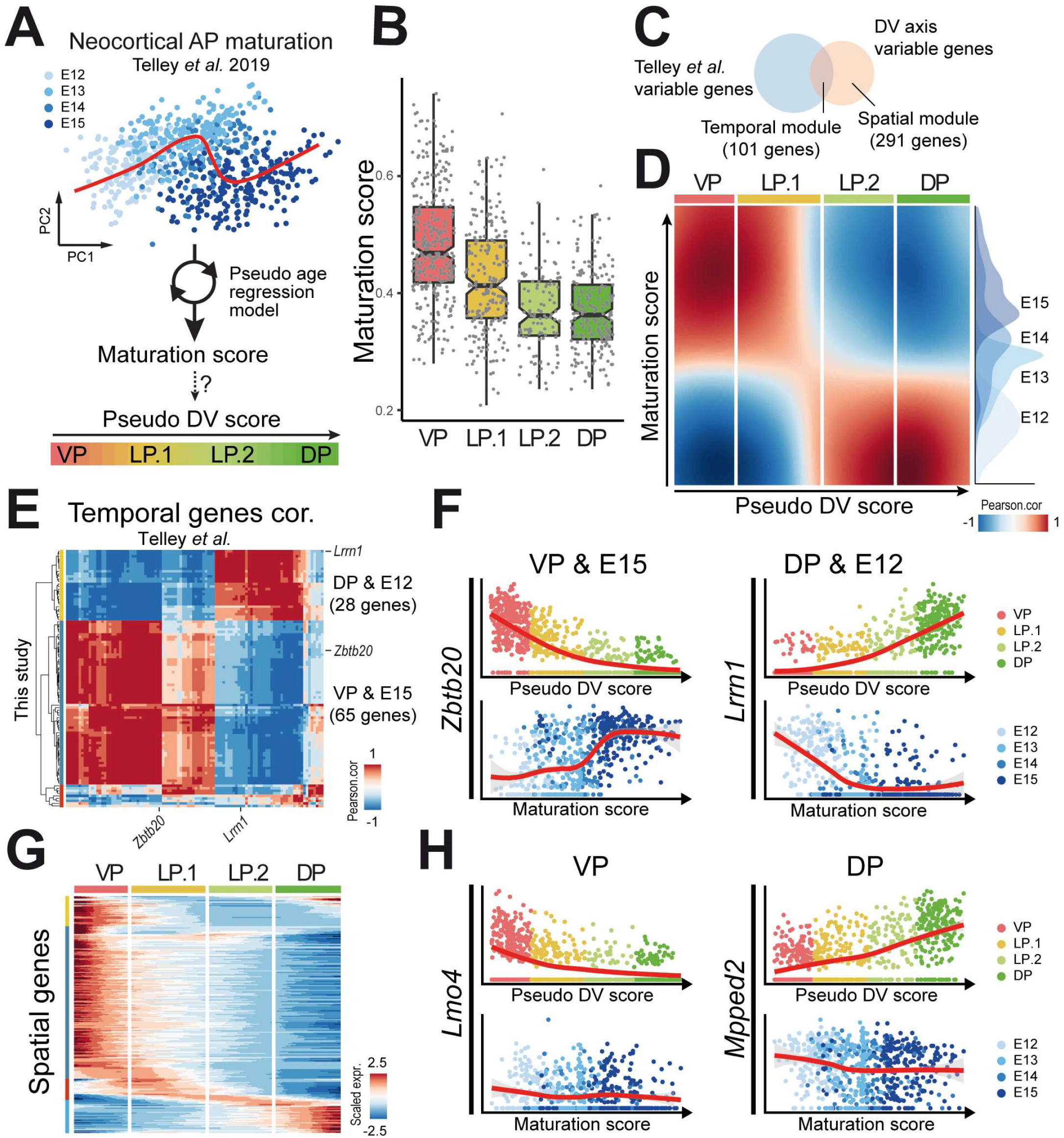
Temporal and spatial contribution to apical progenitors heterogeneity. **A**. A regression model was trained on the AP time series from Telley et al., (2019) and used to predict a maturation score for progenitors in our dataset. **B**. Boxplot depicting the maturation score of APs in each pallial domain. **C**. Venn diagram indicating the overlap between variable genes in the E12-E15 time series of Telley et al. and along the pallial DV axis (this study). **D**. Pearson’s correlation matrix over smoothed expression profiles of temporal module genes showing that maturation and pseudo DV scores are negatively correlated. **E**. Inverse Pearson’s correlation matrix between the smoothed expression profiles of the 101 genes belonging to the temporal module. 66 genes are highly expressed in the E12.5 VP and increase in the DP with time, 35 genes are enriched in the E12.5 DP and decrease with time. **F**. Comparison of representative gene expression trends according to the pseudo DV score in our dataset (top) and maturation score in Telley’s dataset (bottom). **G**. Heatmap showing the smoothed expression of the 291 genes belonging to the spatial module along the pseudo DV axis. **H**. Representative genes expression profile according to the pseudo DV score in our dataset (top) and maturation score in Telley’s dataset (bottom) illustrating time-invariant genes enriched either in the VP or the DP.

We found that among the 392 pallial variable genes in our dataset, 101 also vary during dorsal cortex AP temporal maturation, therefore representing a “temporal module” of pallial AP identity. Conversely, the remaining 291 genes form a “spatial module” that does not show such variation in time during neocortical development (Fig. 6C, Tables S5, S6). We used genes of the temporal module to investigate whether the temporal-associated signature changes continuously along the spatial and temporal axes. To this aim we performed correlation between the smoothed gene expression profiles computed in the two datasets and found two blocks of strong correlation between VP/LP.1 and E15 APs on the one hand, and LP.2/DP and E12 APs on the other hand (Fig. 6D). The distinction between LP.1 and LP.2 appears very sharp when considering genes from the temporal module, i.e. genes whose expression is turned on or off when dorsal cortex APs switch from the production of deeper layer to upper layer neurons. We found that within the temporal module, 66 genes are enriched in the E12.5 VP and upregulated at E15 in the DP as exemplified by the transcription factor *Zbtb20* (Fig. 6F). They can be pictured as a wave of genes progressing dorsally with time. The remaining 35 genes, such as *Lrrn1*, displayed the opposite dynamics, highly expressed in the DP at E12 and turned off at E15 (Fig. 6E,F, Table S5).

When focussing on the spatial module (i.e. genes whose expression in constant from E12 to E15), we found far more genes upregulated in the VP than in the DP (225 vs 41, respectively), indicating that the spatial signature of the VP is more pronounced than that of any other pallial domains (Fig. 6G,H, Table S6).). *Lmo4* and *Mpped2* exemplify such time-invariant genes enriched either in the VP or the DP, respectively (Fig. 6H). By contrast, only 25 genes were found enriched in the LP.1/LP.2, none of which displayed strong specificity or sharp pattern (not illustrated), indicating that APs in these domains are best defined by the combined low expression of VP- or DP-enriched genes (Fig. 6G).

Altogether, these results indicate that differences between progenitors along the DV axis are explained by the superimposition of a maturation-associated gene module observed between E12 and E15 in the DP, together with a spatial stage-independent VP or DP identity.

### Differentiation trajectories can be inferred from AP populations to mature neurons

We then attempted to reconstruct developmental trajectories from AP to LN, assuming that most of the glutamatergic neurons we characterised were collected together with their progenitor of origin. Because CRs are known to migrate tangentially over long distances, our dissections must have not only excluded some VP-derived CRs that migrated away, but also included CR derived from progenitors located outside of the dissected area (hence absent from the dataset). We therefore excluded CR cells from our lineage analyses.

In order to link neuronal populations to their progenitor pools of origin and identify their lineage-specific intermediate states we used FateID (Herman et al., 2018). Briefly, FateID algorithm starts from the most differentiated clusters (as defined in Fig. 2D) and progresses backward iteratively using a random forest classifier to compute a fate probability bias for each cell. From the seven mature populations that were initially seeded, only the two most distant trajectories, leading to *Bhlhe22*^+^ and *Nr4a2*^+^ populations, could be confidently reconstructed back to the AP state (Figs 7A,B, S7A). For the remaining neuronal types, lineage-specific transcriptional signatures only emerged at postmitotic stages suggesting that fate acquisition does not occur at the same differentiation step in all lineages or is not strongly transcriptomically encoded and remains hidden to scRNAseq. Alternatively, we cannot formally exclude the possibility that some lineage intermediates were missed in the dissection due to cell migration (especially for the *Foxp2*/*Lhx5*^+^ population for which the FateID pipeline was unable to reconstruct even the beginning of a trajectory).

**Figure 7.**
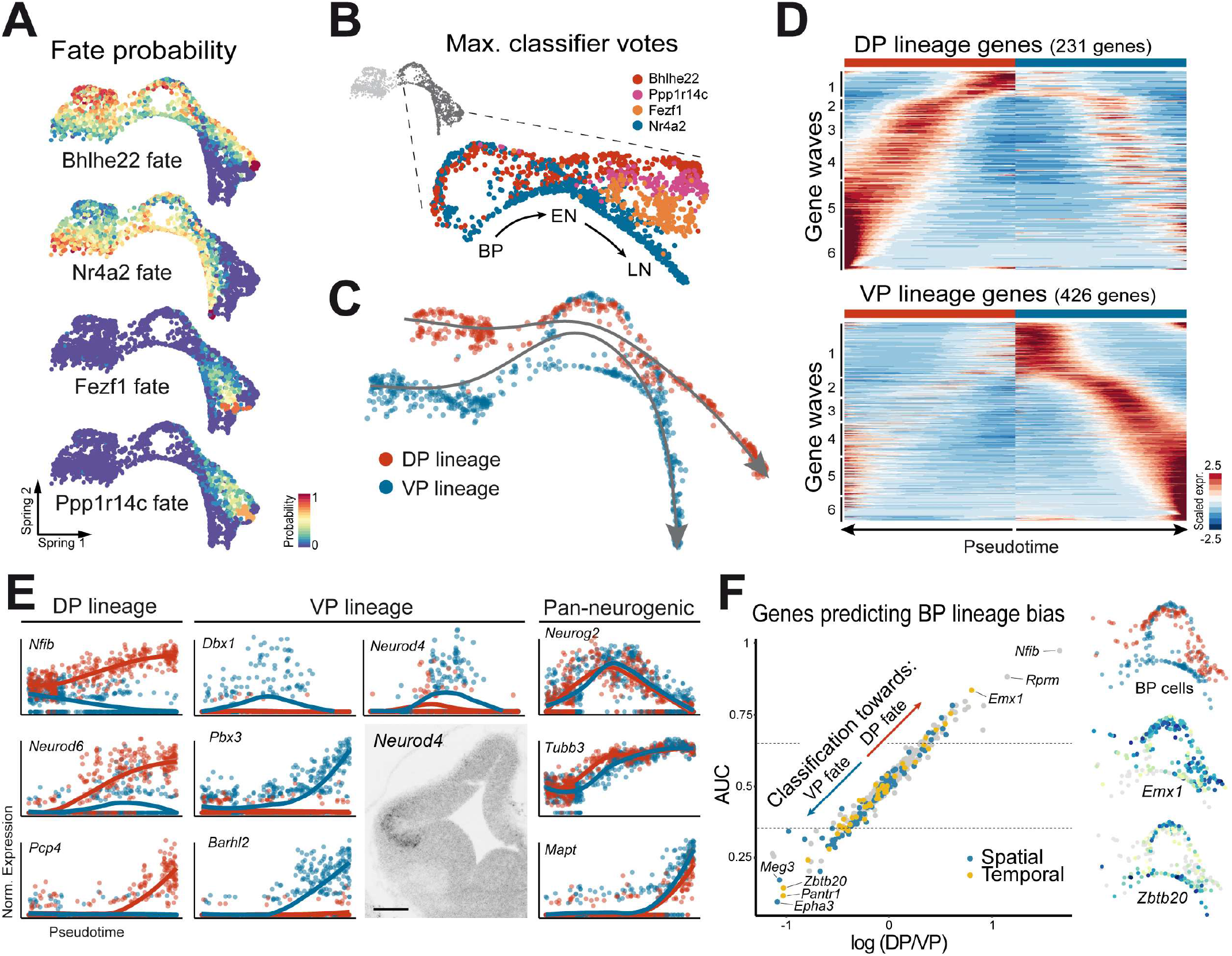
VP and DP lineage reconstruction. **A**. SPRING visualization of the fate probalility predicited by FateID, expressed as the percentage of vote received from the random forest classifer, towards *Bhlhe22*+, *Nr4a2*+, *Fezf1*+ or *Ppp1r14c*+ neurons. **B**. Expanded view of cells at the transition state coloured according to the fate with strongest predicted bias, indicating differences in the rise of transcriptional signature across lineages. **C**. SPRING plot visualization of the fitted principal curves over cells that were confidently assigned to the VP or DP lineage. **D**. Expression profile of genes differentially regulated along the DP or VP trajectories and clustered based on their dynamics along the pseudotime axis. **E**. Comparison of selected gene expression along pseudotime in the DP (red) or VP (blue) trajectory. In situ hybridisation for *Neurod4* on E12.5 coronal section showing its enrichment in the VP/LP intermediate zone. The overlapping dynamics of pan-neurogenic genes validates the pseudotime alignment of the two trajectories. Scale bar: 200 µm. **F**. Scatter plot of the Area Under the Curve (AUC) of the Reciever Operating Characteristic (ROC) measuring individual genes performance to classify BP cells as belonging to VP or DP lineages. Genes of the temporal and spatial modules are indicated in blue and yellow, respectively. SPRING plot visualisation of the expression of *Emx1* and *Zbtb20* in BP cells.

Nevertheless, classification scores for AP obtained from this approach supported histological observations, by giving higher *Nr4a2*^+^ neurons bias to the VP cluster while giving higher *Bhlhe22*^+^ neurons probability to the DP cluster (Fig. 7A,B,C). Interestingly, we observed a clear distinction between the two lineages within cells at the BP progenitor state (Fig. 7B,C) with most of the cells undergoing direct neurogenesis being attributed to the VP/Nr4a2 lineage. Although we believe this should be taken with care due to the slightly higher sampling of the VP trajectory, our data suggest that lineage amplification through BP cycling is not the same for all cell types.

We identified 231 genes significantly upregulated along the DP trajectory and 426 along the VP trajectory (Fig. 7C,D and Tables S7, S8). Those were found at all steps of differentiation even though signature specificity increased with pseudotime (selected examples are shown in Figs 7E, S7B), consistent with the idea that neuronal identity is acquired progressively throughout differentiation. We validated the approach with the observation that *Dbx1*^+^ cells are almost exclusively found in the VP-derived Nr4a2 lineage. Furthermore, we identified additional genes whose expression peaked at the BP stage in the VP trajectory, such as *Neurod4, Dleu7* or *Svil*, but failed to identify counterparts in the DP lineage (Figs 7E, S7B). Consistently, in situ hybridisation confirmed that *Neurod4* expression is highly enriched in the VP subventricular zone with respect to the DP. In the DP lineage, genes such as *Nfib, Neurod2/6* or *Pcp4* are sequentially upregulated (Fig. 7E). By contrast pan-neurogenic genes such as *Neurog2, Tubb3* or *Mapt* displayed identical patterns of expression in the two lineages (Fig. 7E) suggesting that neuronal differentiation proceeds at a similar pace in pseudotime, if not in real time. We then attempted to determine to which extent neuronal fate specification relies on information transmitted from radial glia or from the onset of lineage-specific gene expression. To do so, we performed a receiver operating characteristic (ROC) curve analysis and identified the genes whose expression can best classify BPs between VP and DP lineages (Fig. 7F and Table S9). Out of those best predicting the DP lineage bias, only few were already differentially expressed among APs such as *Emx1*. By contrast, the genes best predicting the VP lineage bias in BPs were often found differentially expressed in APs, belonging to both the temporal (e.g. *Zbtb20* and *Pantr1*) and spatial modules (e.g. *Epha3* and *Meg3*) described earlier. These results suggest that commitment to the VP fate mostly relies on genes expressed in APs, with the contribution of both spatial and temporal modules, whereas commitment to the DP fate mostly involves the upregulation of genes that were not already distinguishing APs.

Taken together, our data indicate that significant differences exist between the specification program of distinct lineages. They suggest that neuronal differentiation – i.e. the acquisition of a neuronal phenotype - proceeds independently of fate acquisition. They also point to the idea that, at least for some lineages, cortical progenitors are already strongly-biased at the AP stage. Our work therefore allows, for the first time, to disentangle the sets of genes involved in neuronal differentiation from those controlling neuronal subtypes specification in different regions of the developing cerebral cortex.

## Discussion

In recent years, multiple studies focused on the establishment of neuronal diversity during cortical development (Loo et al., 2019; Mayer et al., 2018; Telley et al., 2016; Telley et al., 2019; Vitali et al., 2018). Because of its tremendous expansion in primates, the neocortex has attracted most of the attention. Conversely, lateral and ventral regions of the cerebral cortex have been neglected despite being involved in key physiological processes. The tetrapartite model of cortical development states that VZ progenitors belong to four consecutive domains (VP, LP, DP and MP). The precise contribution of LP and VP have long been discussed (Medina et al., 2004; Puelles, 2014; Saulnier et al., 2013) and the expression of *Nr4a2* has been instrumental in the recent redefinition of LP and VP derivatives (Puelles, 2014; Puelles et al., 2016a). We have shown that most of the early VP derivatives (as identified by Dbx1 genetic tracing) consist in *Nr4a2*^+^ cells, in addition to some *Foxp2*^+^ cells and p73-negative CR (Bielle et al., 2005; Griveau et al., 2010). This comes as a surprise given that *Nr4a2* is only reported to be expressed in the presumptive LP derivatives that are the claustrum and dorsal endopiriform nucleus (Puelles et al., 2016a; Watson and Puelles, 2017), to which the Dbx1 lineage is not supposed to contribute (Puelles et al., 2016b). By contrast, the *Fezf1*^*+*^ population we characterised is most likely derived from the LP according to its position in an Emx1-derived domain dorsal to Nr4a2 cells, and possibly corresponds to neurons that will eventually settle in the lateral amygdala, as expression of *Fezf1*^*+*^ has been reported in this region (Hirata et al., 2006; Kurrasch et al., 2007). This population would represent an early-born LP contribution to the amygdala, as previously postulated (Cocas et al., 2009; Gorski et al., 2002). Our findings therefore point to the idea that LP and VP derived structures are perhaps not as sharply distinguished as initially anticipated (Puelles et al., 2000).

We captured distinct CR subtypes in our dataset. Because CRs originate from at least 4 regions of the developing forebrain: the septum, hem, VP and thalamic eminence (Bielle et al., 2005; Ruiz-Reig et al., 2017; Takiguchi-Hayashi, 2004) and because these regions are distant from each other and consist in vastly different micro-environments, the question of CR diversity has long been standing. Although a hallmark of CRs is their absence of *Foxg1* expression, we were unable to identify a single gene specifically expressed by all CR subtypes, as exemplified by *Reln* and *Calb2* (Calretinin) which are found in the dorsal cortex preplate as well as in some Foxp2 populations. Although previous work revealed subset-specific features (Griveau et al., 2010; Hanashima et al., 2007), the extent of similarity/differences between CR populations remained largely uncharted. We report that the first and main difference between CR subtypes corresponds to their lateral or medial origin. Interestingly, VP-derived CRs mostly differ from hem, septum and thalamic eminence CRs by lacking a complete set of genes classically used as CR markers (e.g. *Trp73, Lhx1, Ebf3, Cacna2d2*). By contrast, we could not find genes very specifically expressed by VP-derived CRs, suggesting they represent a default fate, consistent with previous studies indicating that only VP-derived CR are produced during conditional inactivation of *Foxg1* (Hanashima et al., 2007). Accordingly, the production of medial CRs would follow a specific trajectory leading to the expression of the aforementioned gene module. CRs are highly motile cells and display complex migratory patterns resulting in a certain degree of intermingling between the different populations and dispersion over large territories (Barber et al., 2015; Griveau et al., 2010; Yoshida et al., 2006). We found out that the two clusters of medial CRs (*Trp73*^+^) can be distinguished by the opposite expression of gene modules correlating with spatial positioning of neurons along the DV axis, consistent with the hypothesis that part of CR identity is imposed by the environment in which they settle. Such a spatial component to neuronal identity has also recently been highlighted for pyramidal neurons of the neocortex (Yao and Tasic, 2020).

Sharp segregation of progenitor domains in the developing spinal cord has long suggested that spatial patterning is a key driver of neuronal diversity in the central nervous system (CNS). However, temporal patterning was recently proposed to be also at play and to involve the same gene networks throughout the CNS (Sagner et al., 2020). In the neocortex, the graded expression of patterning genes, the sequential production of deep and upper layers neurons and the changes in the transcriptomic signature associated with AP maturation (Okamoto et al., 2016; Telley et al., 2019) clearly point to the importance of temporal mechanisms. However, the spatial component has been systematically underestimated by recent studies implementing scRNAseq approaches, perhaps simply because the prospective neocortex (dorsal pallium) focused most of the attention at the expense of the ventral and lateral pallium. Nevertheless, one should keep in mind that spatial and temporal regulations are intimately linked. This is perhaps best exemplified by the neurogenic gradient that leads to the generation of ventral neurons earlier than dorsal ones and is associated to opposite gene waves progressing along the DV axis of the brain. The “temporal module” we describe consists in genes whose expression in neocortical progenitors not only changes with time but is also spatially restricted at E12.5 (mostly to the VP). Similarly, the membrane potential of cortical APs, which was shown to change with temporal maturation, is also different between dorsal and ventral cells at the same developmental stage (Vitali et al., 2018). In the future, it would be interesting to investigate within the DP how much of the variability among APs at a given stage can be attributed to the position cells occupy on the DV axis, and to which extent such differences rely on the temporal maturation gradient along this same axis. In addition, it remains to be established whether spatial and temporal associated gene regulatory networks are independent or cross regulated. In an evolutionary perspective, a switch form spatial to temporal patterning, involving the co-option of an early VP-specific gene module by DP progenitors, has been proposed to be an important innovation for the acquisition of upper layers in mammals (Luzzati, 2015). Consistently, the loss of function of *Zbtb20* or *Tfap2c*, which expression are first restricted to the VP before progressively expanding in dorsal progenitors, result in upper layer neurons generation defects (Pinto et al., 2009; Tonchev et al., 2016). It will thus be interesting in future studies to compare gene expression in cortical progenitors across different species and investigate the extent of conservation, in the composition and dynamics, of the spatial and temporal gene modules.

When looking at the identity of APs along the DV axis of the pallium (this study) or through time (Telley et al., 2019), progenitors appear relatively smoothly distributed with no obvious sharp transition between them. This raises the question of how we should define progenitor domains when the distinction between APs is continuous. It is interesting that a similar question was raised by Wulliman (Wullimann, 2017), based on purely neuroanatomical grounds. The lack of strong experimental arguments supporting the existence of fate-restricted progenitors in the DP argues in favour of a model whereby what we currently define as domains do not exist *per se* but rather that a continuity of APs are biased towards alternative fates and that the probability to tip over one fate or another changes with the DV position, according to the graded expression of instructive factors.

We found that APs occupying the most extreme position in the gradient (VP and DP) display the strongest spatial signature and bias towards specific terminal fates. By contrast, APs localised at intermediate positions along the DV axis do not show such a strong transcriptional identity that could be transmitted to BPs, explaining why our lineages reconstruction were only effective when comparing the most distant terminal fates in our dataset. However, the near-perfect tangential segregation of *Ppp1r14c*^+^ and *Fezf1*^+^ neurons argues in favor of an equivalent spatial segregation of their respective progenitors in the LP. Therefore, we propose that at the cell population level, bias towards a specific lineage progressively decreases, as progenitors are located closer to a transition zone between two domains while the opposite probability towards generating the neighbouring cell type would increase (Fig. 8A). Such a model is supported by the observation that genetic tracing using sharp domain-restricted enhancers results in fuzzy fate maps with graded contribution to adjacent cortical structures (Pattabiraman et al., 2014). At the individual cell level, some progenitors would adopt a deterministic commitment towards one lineage while others, located at transition zones, would undergo a succession of probabilistic choices between two adjacent fates. Alternatively, progenitors commitment could be resolved very early on, when the cortical primordium is being patterned, resulting in transition zone containing unsorted APs each bearing distinct pallial domain fate potential. Clonal fate tracing techniques provide a powerful experimental paradigm to distinguish between the two alternatives (Gao et al., 2014; He et al., 2012; Llorca et al., 2019) and we expect future studies taking advantage of such approaches to clarify the link between the behaviour of individual progenitors and their position on the DV axis.

**Figure 8.**
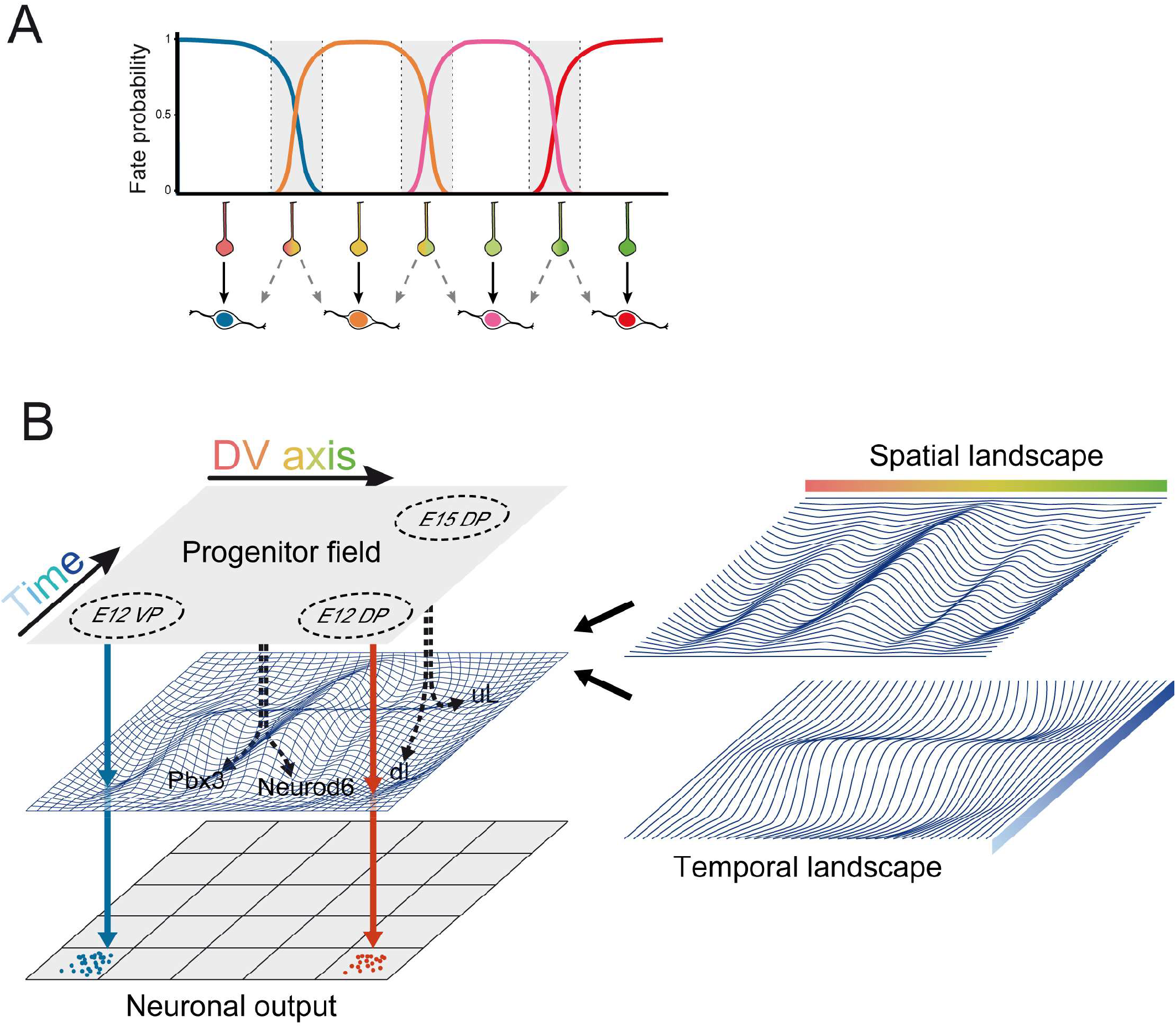
Model of superimposition of spatial and temporal landscapes. **A**. Shematic representation of the probability to specify different neuronal identities according to the progenitor position along the dorso-ventral axis. While progenitors located in the middle of one domain will give rise to the same neuronal fate in a deterministic manner, progenitors sitting at the transition between two adjacent domains (grey area) might undergo a succession of probabilistic decisions between two alternative fates. **B**. A progenitor field represents APs according to their temporal maturation and location along the DV axis. The superimposition of temporal and spatial landscapes dictates the discrete neuronal fate towards which differentiating cells are canalised. Solid arrows depict the differentiation trajectory followed by progenitors that were initially strongly biased. Dotted arrows represent the differentiation of progenitors located at transition zones and for which random fluctuations in the transcriptional state will have more influence on the choice between alternative fates. This is the case between Neurod6 and Pbx3 lineages along the DV axis, or at the transition between upper layer (uL) and deep layer (dL) neurons along the time axis.

The continuity between APs contrasts with the discrete nature of LN, raising the question of the time point at which cell fate is assigned. In the neocortex, graft experiments of DP progenitors indicated that APs display greater plasticity than BPs which are already committed to specific neuronal fates (Oberst et al., 2019). In our dataset, the bias for *Bhlhe22*^+^ and *Nr4a2*^+^ lineages is fairly constant from AP to LN and does not differ between AP and BP, strongly supporting an early commitment taking place at the AP level for these lineages. Yet, the fact that not all trajectories were reconstructed back to APs could also indicate that fate restriction occurs at distinct steps in each lineage, although we cannot formally exclude that commitment indeed occurs at the AP level but remains hidden at the transcriptomic level as shown in other developmental systems (Weinreb et al., 2020; Ziffra et al., 2020). Nevertheless, our data indicate that major transcriptional differences between lineages rise after cells exit the BP state.

In an attempt to summarise our findings we adapted the classical Waddington metaphor to represent the influence of both space and time on the same landscape (Fig. 8B). In this representation, AP identity is encoded in its coordinates on a spatio-temporal map. Upon exit from a progenitor state, this coordinate sets the initial position of the cell onto the developing landscape which results from the superimposition of spatial and temporal components. In this model, cells starting from progenitor fields at extremities of the landscape will systematically fall into a basin of attraction which will canalise their differentiation towards the same attractor state (i.e. neuronal fate). By contrast, APs close to the tipping point separating two basins of attraction, will be more sensitive to small random fluctuations in their initial state, which might drive them towards either one of two alternative fates. Changes in the regulatory dynamic of genes that underlie the topology of spatial and temporal landscapes result in the modification of the amount and identity of neuronal types being produced from a specific domain. Accordingly, one can also speculate that evolutionary modification of these landscapes might give access to previously unexplored attractors and therefore allow to generate novel neuronal types.

## Acknowledgements

The authors wish to acknowledge Aurelia Dujardin, Camelia Piat and the Animalliance platform of the *Imagine* Institute for animal care; the SFR Necker imaging and histology platforms for help with acquisition; Fanny Coulpier from the IBENS genomics platform for sequencing; Christine Bole and Aurore Pouliet from the *Imagine* Institute genomics platform for helpful discussions; Frédéric Tores, Jean-Marc Plaza and Patrick Nitschke from the *Imagine* Institute bioinformatics platform for implementing the offline version of SPRING Viewer; 10X Genomics and Miltenyi Biotec technical support for their help in the implementation of scRNAseq. We thank all members of the Pierani lab, Marine Luka and members of the Ménager lab as well as members of the Rausell lab for helpful discussions. We are grateful to Anne Teissier and Oriane Blanquie for critical reading of the manuscript. MXM is the recipient of an Allocation Spécifique PhD fellowship from the École Normale Supérieure, FC is an Inserm researcher, AP is a CNRS investigator. This work was supported by grants from Agence Nationale de la Recherche (ANR-15-CE16-0003-01) and Fondation pour la recherche médicale (Équipe FRM DEQ20130326521) to AP and State funding from the Agence Nationale de la Recherche under “Investissements d’avenir” program (ANR-10-IAHU-01) to the *Imagine* Institute.

## Authors contribution

Conceptualization: AP, FC and MXM

Methodology: FC and MXM

Software: MXM

Validation: FC and MXM

Formal analysis: MXM

Investigation: AWC, FC, MXM and YS

Data Curation: FC and MXM

Writing – original draft preparation: FC and MXM

Writing – review and editing: AP, FC and MXM

Visualization: FC and MXM

Supervision: AP and FC

Project administration: AP and FC

Funding acquisition: AP

## Competing interests

No competing interests declared.

**Supplementary Figure 1.**
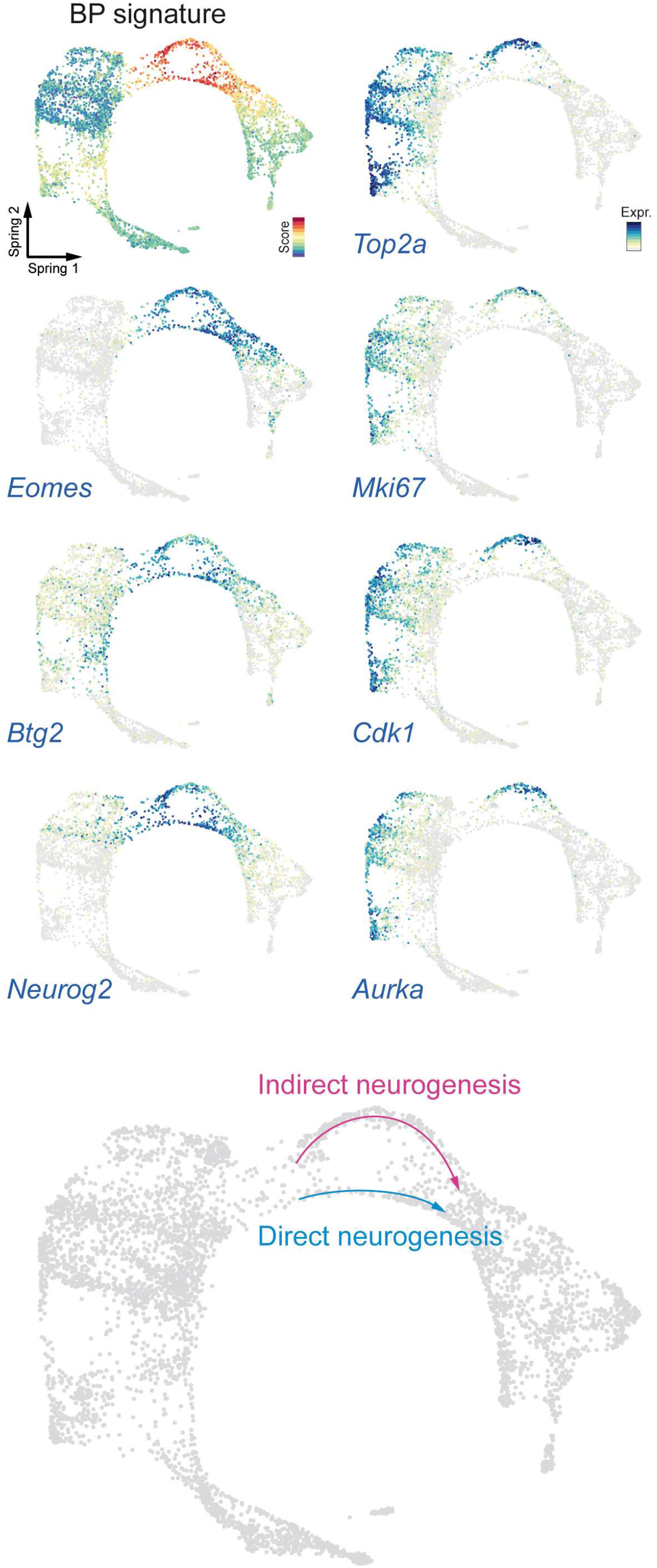
Direct and indirect neurogenesis. SPRING visualisation of the BP signature score as well as the expression of BP genes *Eomes* (Tbr2), *Btg2* (Tis21) and *Neurog2*. Proliferation genes such as *Top2a, Mki67, Cdk1* and *Aurka* are only expressed by a fraction of BPs, leading to the segregation of two trajectories corresponding to direct and indirect neurogenesis.

**Supplementary Figure 2.**
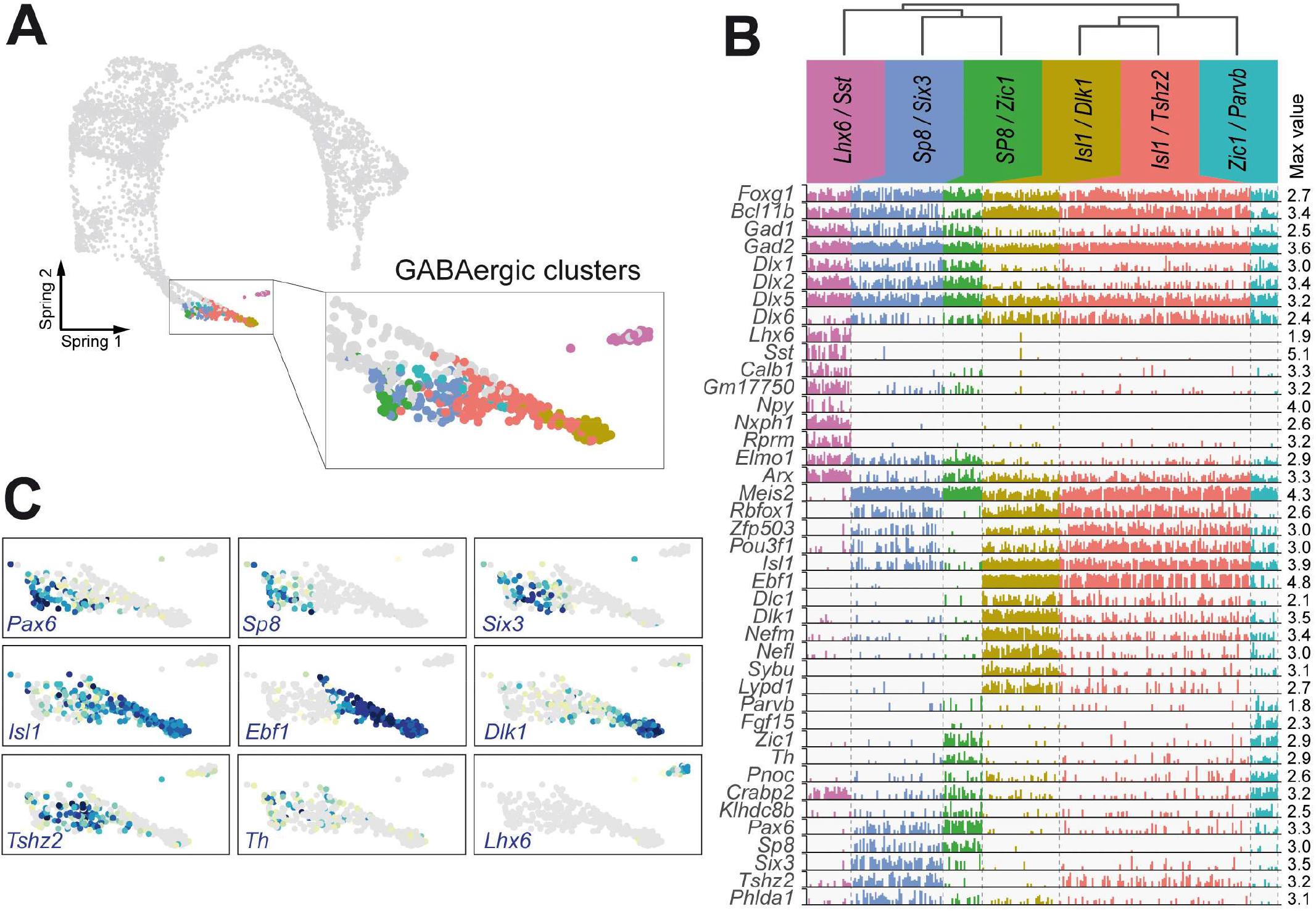
Diversity of GABAergic neurons. **A**. SPRING representation of the dataset illustrating the 6 clusters of inhibitory neurons. **B**. Bar plot representing the expression level of selected genes that are differentially expressed between clusters and allow to finely define them. Each bar corresponds to a single cell, bars height corresponds to the expression level of a given gene in a given cell. The dendrogram on the top indicates the hierarchical relation between cell types. **C**. Expression level of selected genes differentially expressed between inhibitory subtypes.

**Supplementary Figure 3.**
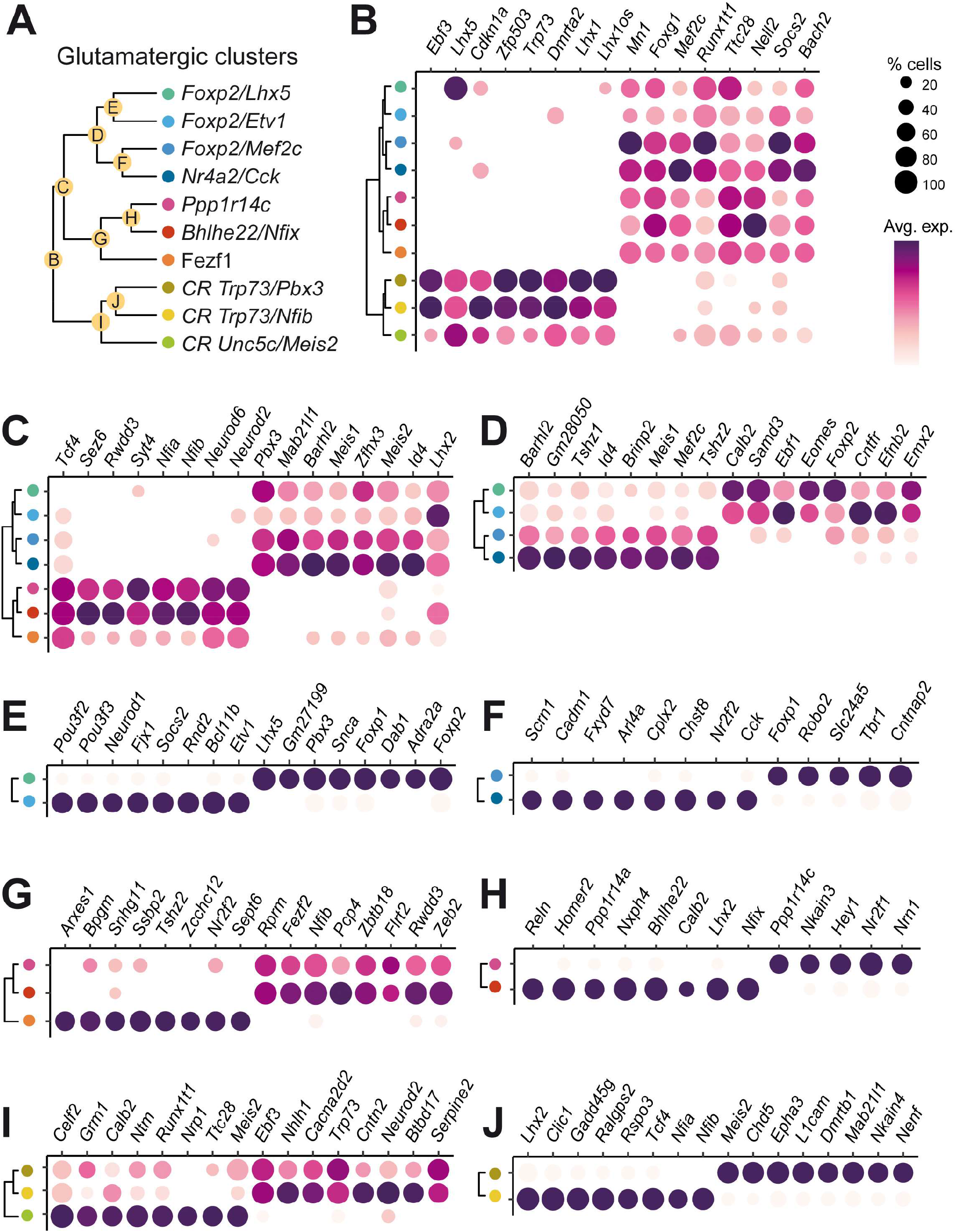
Distinction between excitatory neuron types and classes. **A**. Dendrogram indicating the hierarchical relation between clusters of excitatory neurons. The letter indicated in each node refers to the panel comparing the two branches of the node. **B-J**. Bubble charts comparing the neuronal subtype belonging to each branch of a given node. The size of dots corresponds to the fraction of cells expressing the gene considered, whereas the average expression level is color-coded. Represented genes were selected based on their performance in classifying cells between the two branches considered (AUC-ROC).

**Supplementary Figure 4.**
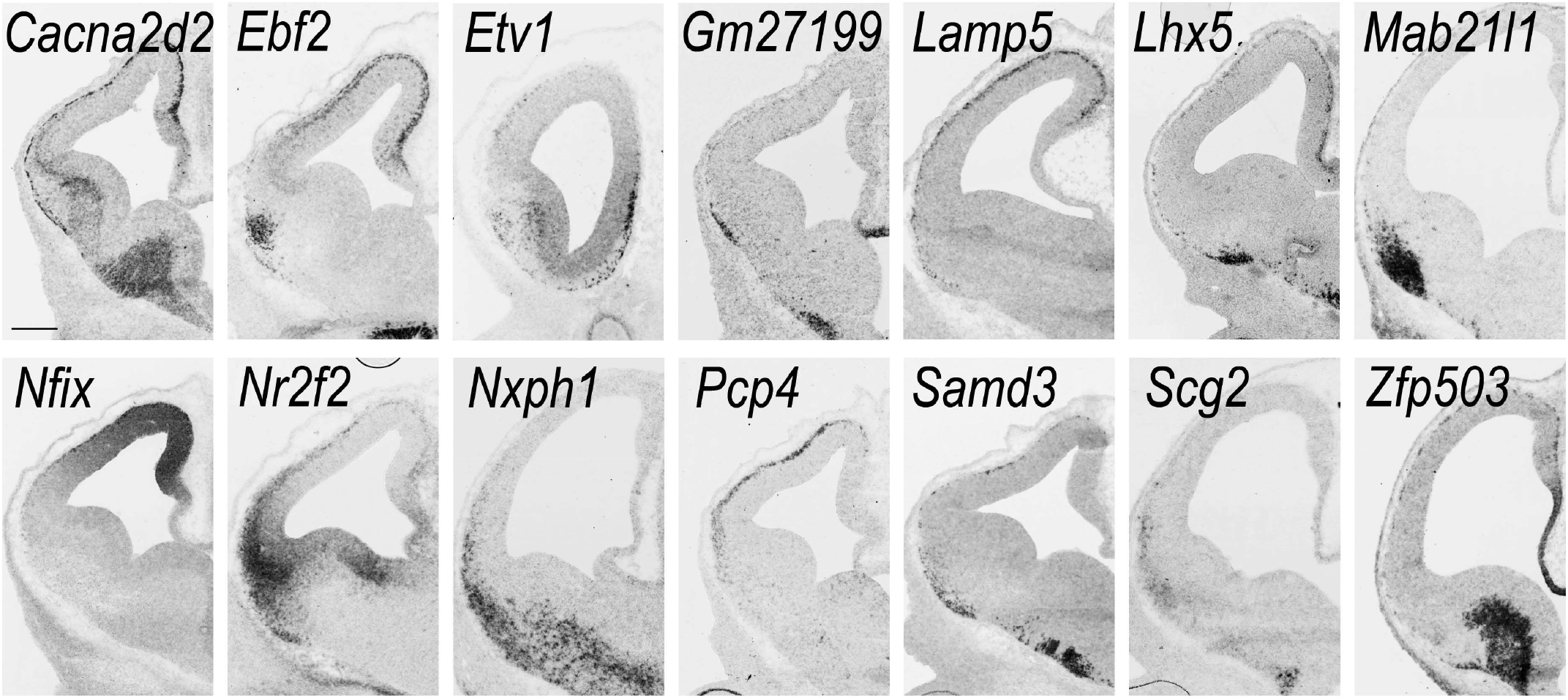
Additional expression patterns. In situ hybridisation of genes differentially expressed in excitatory neurons on E12.5 telencephalon coronal sections. Scale bar: 200 µm.

**Supplementary Figure 5.**
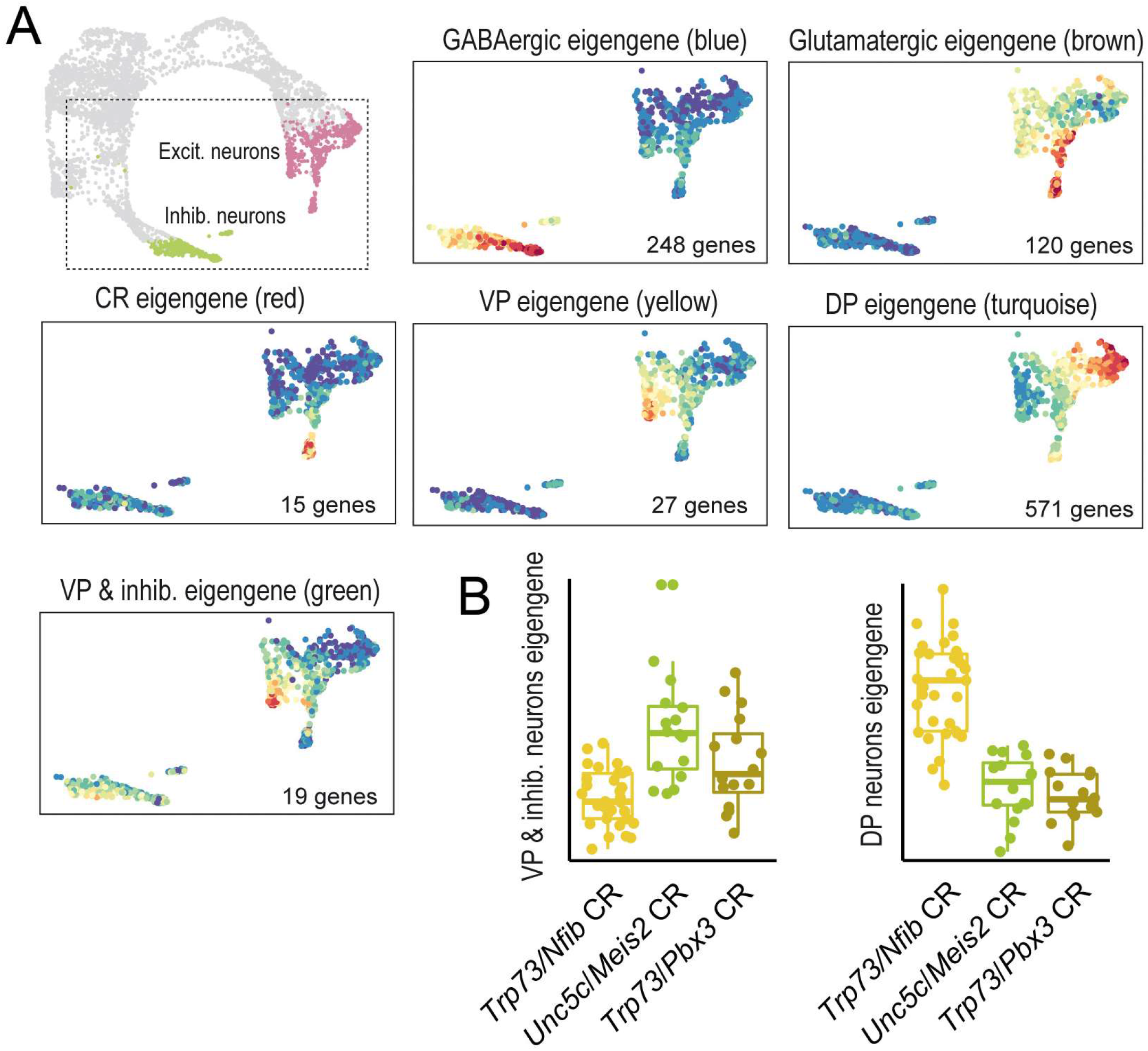
WGCNA analysis. **A**. SPRING representation of all mature neurons present in the dataset that were subjected to WGCNA analysis. The expression profile of the 6 gene coexpression modules identified is color coded from blue (low) to red (high). The colour indicated in between brackets refers to the name of the module in Table S3. **B**. Boxplot illustrating the two eigengenes that segregate CR clusters according to their position in dorsal (*Trp73*/*Nfib* CR) or ventral territories (*Unc5c/Meis2* and *Trp73/Pbx3* CR).

**Supplementary Figure 6.**
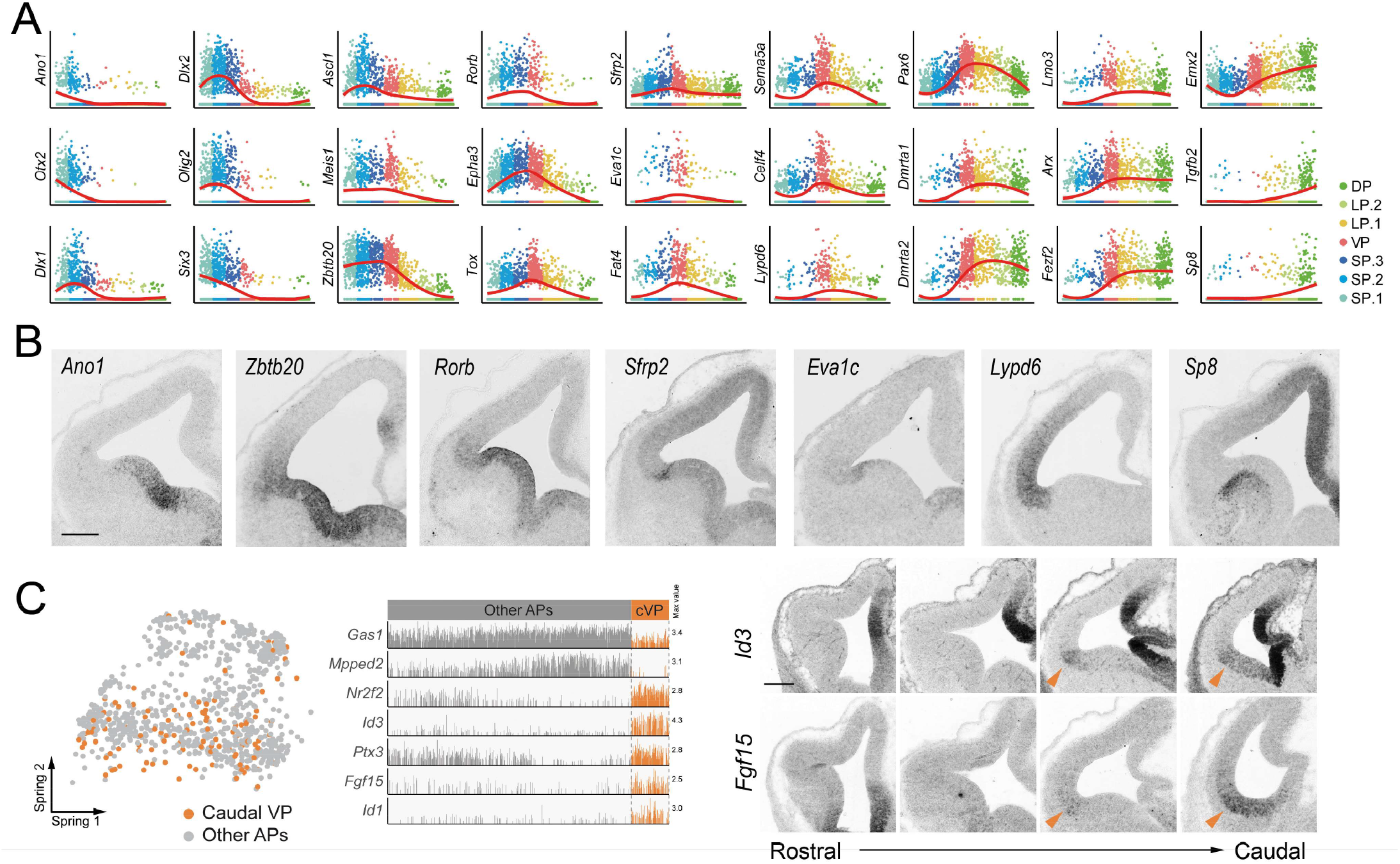
Additional genes with variable expression in APs along the DV axis. **A**. Representative gene expression profiles along the pseudo DV axis. **B**. In situ hybridisation for genes selected among those shown in A. **C**. The SPRING plot on the left highlights caudal VP progenitors identified by iterative graph-based clustering after regressing sources of variation correlating with the pseudo DV position or cell cycle score. The bar plot indicates genes differentially expressed between caudal VP and other progenitors. In situ hybridisation panels on the right exemplify the expression pattern of caudal VP markers along the antero-posterior axis. Scale bar: 200 µm.

**Supplementary Figure 7.**
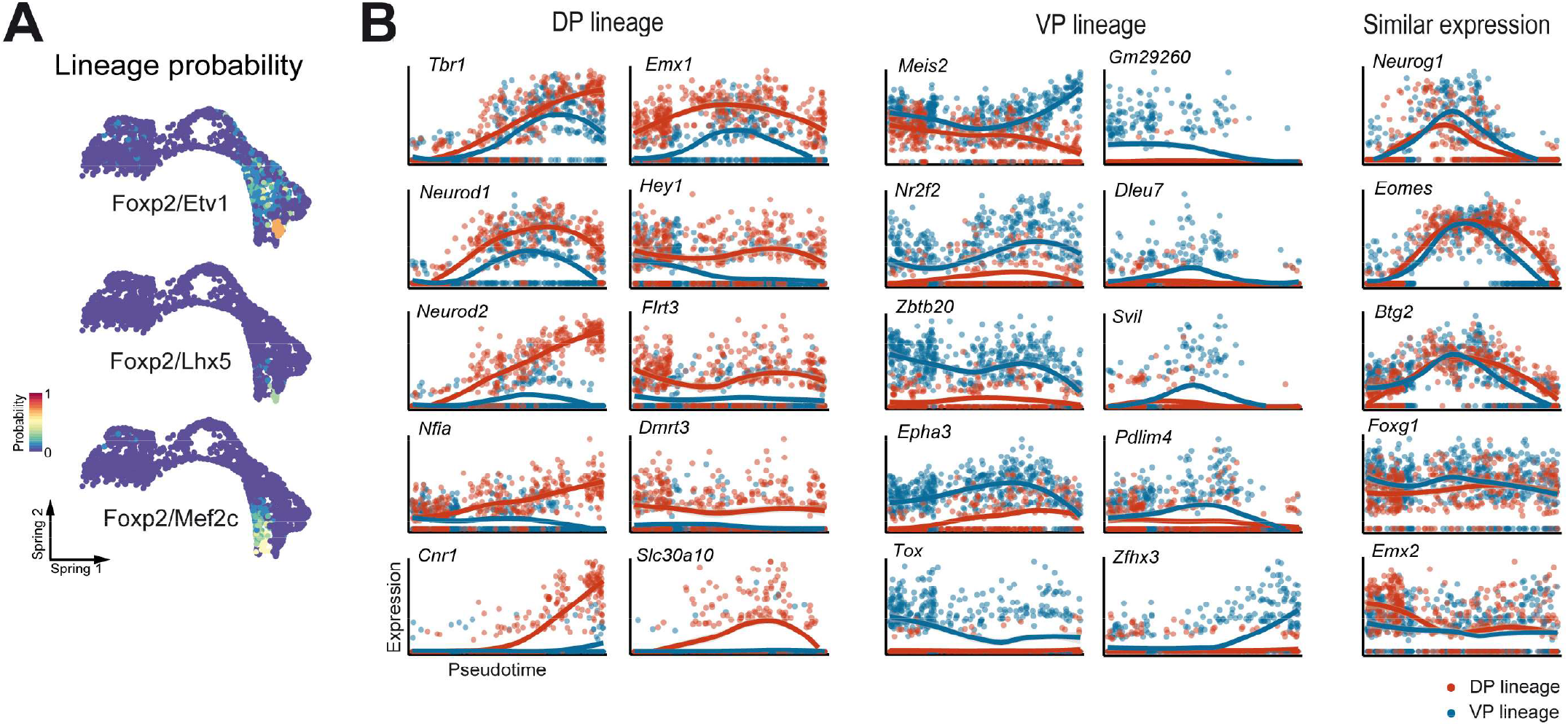
Additional lineage reconstructions. **A**. SPRING visualisation of the different fate probabilities predicted by FateID, expressed as the percentage of votes received from the random forest classifier. None of the cell types illustrated here could be traced back to the level of apical progenitors. **B**. Comparison of representative genes expression along pseudotime in the DP (red) or VP (blue) trajectory.

